# CiDRE^+^ M2c macrophages hijacked by SARS-CoV-2 cause COVID-19 severity

**DOI:** 10.1101/2022.09.30.510331

**Authors:** Yuichi Mitsui, Tatsuya Suzuki, Kanako Kuniyoshi, Jun Inamo, Kensuke Yamaguchi, Mariko Komuro, Junya Watanabe, Mio Edamoto, Songling Li, Tsukasa Kouno, Seiya Oba, Tadashi Hosoya, Shohei Koyama, Nobuo Sakaguchi, Daron M. Standley, Jay W. Shin, Shizuo Akira, Shinsuke Yasuda, Yasunari Miyazaki, Yuta Kochi, Atsushi Kumanogoh, Toru Okamoto, Takashi Satoh

## Abstract

Infection of the lungs with severe acute respiratory syndrome coronavirus 2 (SARS-CoV-2) via the angiotensin I converting enzyme 2 (ACE2) receptor induces a type of systemic inflammation known as a cytokine storm. However, the precise mechanisms involved in severe coronavirus disease 2019 (COVID-19) pneumonia are unknown. Here, we show that interleukin-10 (IL-10) changed normal alveolar macrophages into ACE2-expressing M2c-type macrophages that functioned as spreading vectors for SARS-CoV-2 infection. The depletion of alveolar macrophages and blockade of IL-10 attenuated SARS-CoV-2 pathogenicity. Furthermore, genome-wide association and quantitative trait locus analyses identified novel mRNA transcripts in human patients, COVID-19 infectivity enhancing dual receptor (CiDRE), which has unique synergistic effects within the IL-10-ACE2 system in M2c-type macrophages. Our results demonstrate that alveolar macrophages stimulated by IL-10 are key players in severe COVID-19. Collectively, CiDRE expression levels are potential risk factors that predict COVID-19 severity, and CiDRE inhibitors might be useful as COVID-19 therapies.

**Graphical abstract:** 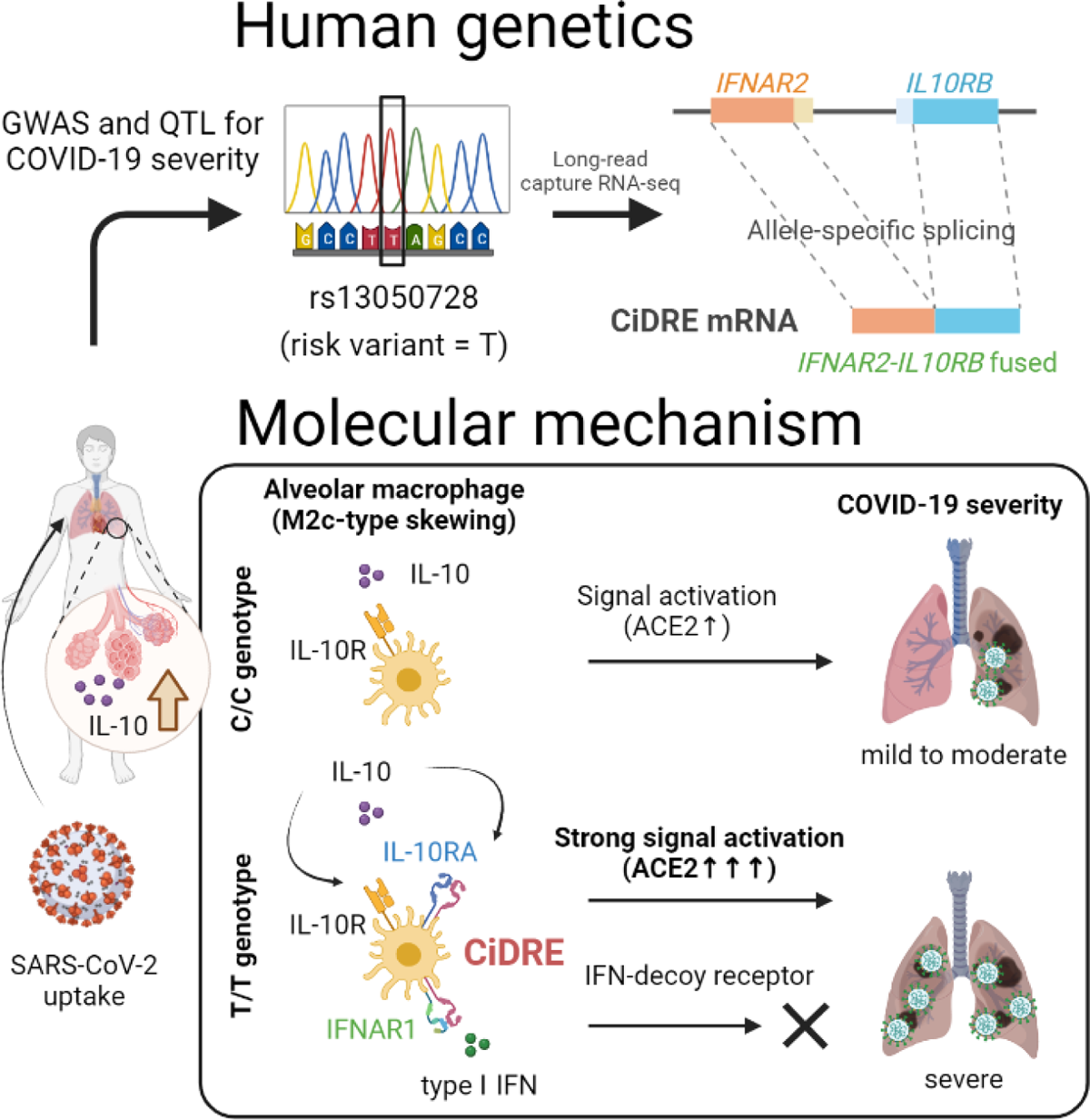

## Introduction

SARS-CoV-2 emerged in December 2019 and led to the COVID-19 pandemic, which has globally disrupted society (*1*). The excess mortality rate of COVID-19 was reported to be 120.3 deaths per 100,000 persons (*2*). The main cause of death is respiratory failure related to severe pneumonia, also termed acute respiratory distress syndrome (*3*). Although there are asymptomatic cases of SARS-CoV-2 infection, some patients progress rapidly into a severe and fatal illness. In a recent retrospective study, several indexes, including sex, age, and body mass index, were reported to be risk factors for the exacerbation of COVID-19 (*4, 5*). However, the detailed molecular mechanism of severe pneumonia in COVID-19 remains unknown. The marked amplification of inflammatory responses triggered by SARS-CoV-2 infection, termed a “cytokine storm”, is considered to be a key process in the course of the disease’s progression and leads to multiple organ dysfunction (*6–8*). Various cytokines (including IL-6) and chemokines produced by virus-infected cells in the lungs are thought to be important for the establishment of cytokine storms. In addition, the accumulation of several types of immune cells, including monocytes, dendritic cells, and CD4 and CD8 T-cells, by chemotaxis facilitates this process. In the first step of this mechanism, SARS-CoV-2 invades host cells via the spike (S) glycoprotein that normally binds to the host cell-surface receptor, ACE2 (*9–12*), which is expressed abundantly in proximal airway cells of the human respiratory system and partially in the distal epithelium, including on alveolar epithelial cells (*13, 14*). Because the focus of cytokine storms is the alveolar space, it is unclear how proximal airway-dominant infections of SARS-CoV-2 lead to widespread inflammation in the distal alveolar area. In alveoli, alveolar macrophages, mainly involved in the maintenance of the immune system, are considered low-ACE2-expressing cells. However, recent research has suggested that alveolar macrophages are susceptible to SARS-CoV-2 infection and are involved in the virus-induced cytokine storm (*15, 16*). In this study, we focused on alveolar macrophages and investigated their functional involvement in COVID-19 severity.

### IL-10 is critical for COVID-19 severity

To investigate the histological changes caused by infection with SARS-CoV-2 in a hamster model, we stained samples of lungs infected with the virus. Histological evaluation of the infected lungs indicated the progression of autophagy and apoptosis but not fibrosis (**Fig. 1A**). Next, we investigated the comprehensive gene expression pattern differences between healthy lungs (Mock group) and SARS-CoV-2-infected lungs (COVID-19 group) using RNA sequencing (RNA-seq) and extracted 1624 upregulated differentially expressed genes (DEGs) and 1427 downregulated DEGs (**Fig. 1, B and C, and fig. S1A**). In addition, we performed enrichment analysis using gene ontology (GO) and extracted cellular processes enriched for the upregulated DEGs (**table S1 and S2**). A Venn diagram was created for five GOs expected to involve macrophages, including immune-response-related, cell death, and autophagy signatures, which were selected based on histological analysis (**Fig. 1D**). We identified five genes (*Psen1*, *Ticam1*, *Pycard*, *Trem2*, and *Il10*) that overlapped across all GOs. Among these genes, *Il10* was the most upregulated in the SARS-CoV-2 infection group compared with uninfected normal lungs (**Fig. 1E**). IL-10, a soluble protein with anti-inflammatory properties, was reported to induce the differentiation of macrophages into anti-inflammatory M2c-type macrophages (*17, 18*). However, no reports have investigated the tissue-specific effects of IL-10 on alveolar macrophages *in vivo* during SARS-CoV-2 infection. A histological assessment of the infected lungs showed that SARS-CoV-2 appeared in the central bronchial epithelium 1 day after infection and then spread through the alveolar space within a few days (**fig. S1B**). In addition, our investigation of IL-10 localization revealed that it was produced at high levels in the bronchial epithelium during COVID-19 infection (**Fig. 1F**). These results suggest that IL-10 secreted by the bronchial epithelium infected with SARS-CoV-2 has an influence on the spread of the virus in lung parenchyma and leads to severe inflammation. To investigate the importance of IL-10 in human COVID-19 severity, clinical serum samples from patients with SARS-CoV-2 were collected after hospital admission (early-stage post-infection), and the concentrations of IL-10 and other serum factors were measured. Based on the extent of disease progression after diagnosis, the patients were divided into two groups: Stable and Progressive (**fig. S2A**). Interestingly, a significant difference in serum IL-10 was observed between the progressive and stable group in the early stage of SARS-CoV-2 infection (after admission), whereas there were no significant differences in other inflammatory cytokines, including IL-6, that have been reported to be highly expressed in the late stages of infection (*19*) (**Fig. 1G and fig. S2B**). Consistent with recent clinical research, CRP levels were also significantly different between the two groups (*20*) (**table S3**). Collectively, these data suggest that IL-10 is a critical factor involved in COVID-19 severity in the early stage of SARS-CoV-2 infection in humans.

**Fig. 1.**
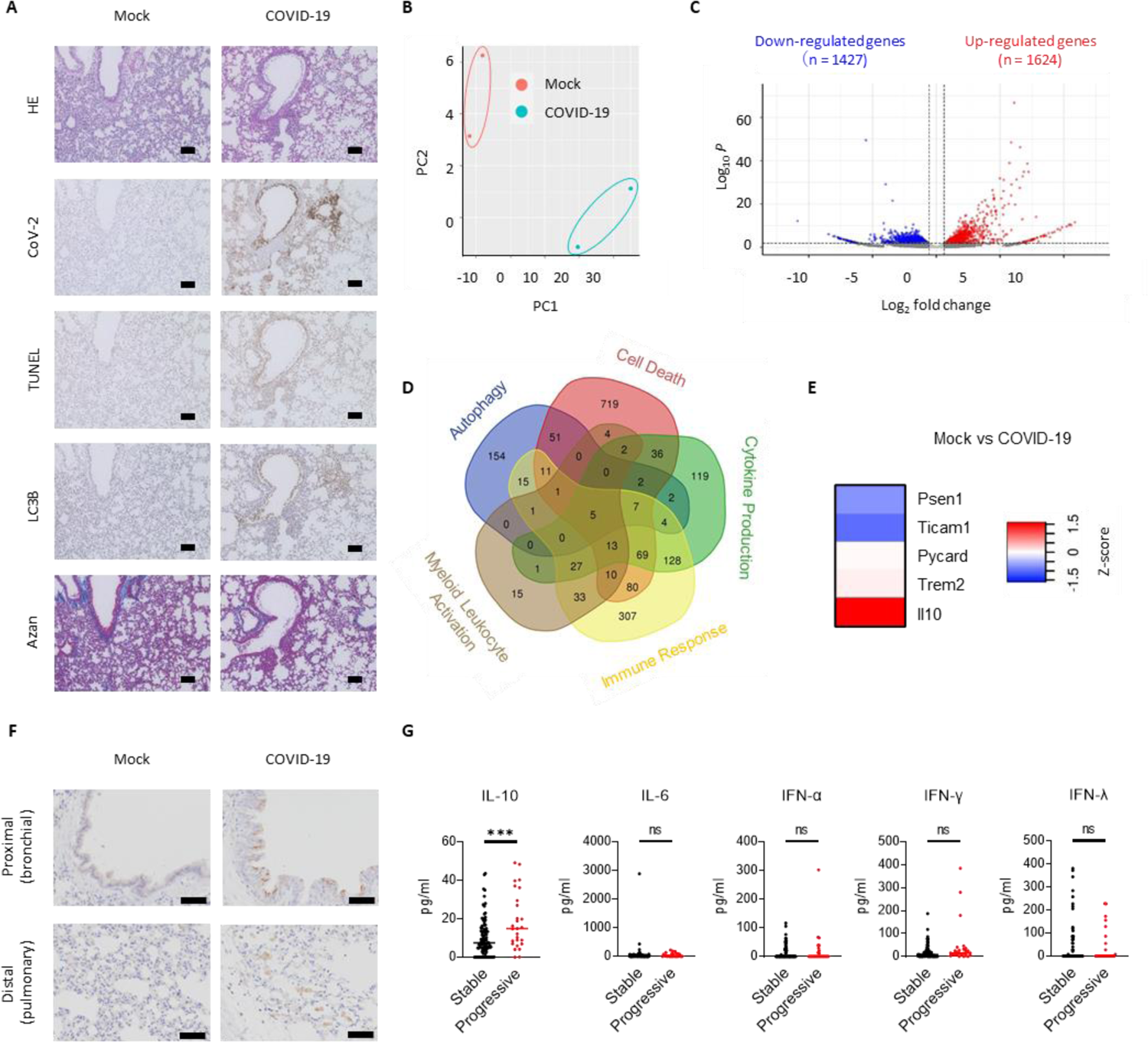
IL-10 is a key regulator of COVID-19 severity. **(A)** Hematoxylin and eosin (HE) staining (top), immunohistochemistry of SARS-CoV-2 (second from the top), TUNEL (third from the top), immunohistochemistry of LC3B (fourth from the top), and Azan (bottom) staining of hamster lung specimens. Scale bars, 100 µm. **(B)** Principal component analysis (PCA) of RNA-seq data between Mock and COVID-19 groups (n=2). **(C)** Volcano plot showing DEGs. The upregulated genes are marked in red and downregulated genes are marked in blue. **(D)** Venn diagram based on five GO groups related to important cellular processes in COVID-19. Each GO includes the upregulated DEGs in COVID-19. **(E)** Heatmap showing the overlapping genes within the five GOs. **(F)** Immunohistochemical staining of IL-10 in lung sections. Scale bars, 50 µm. **(G)** Comparison of serum levels of IL-10, IL-6, IFN-α, IFN-γ, and IFN-λ between two groups of COVID-19 hospitalized patients. The patients were divided into Stable (no disease progression through treatment, n=95) and Progressive groups (disease progression after admission, n = 27). Patient classification criteria are shown in **fig. S2A**. The dots represent individual patients and the error bars indicate the median. Statistical significance was determined by a Mann–Whitney *U-*test.

### M2c-skewing is a key cellular process in SARS-CoV-2 susceptibility of alveolar macrophages

To clarify the role of high levels of IL-10 in severe COVID-19 patients, we investigated a potential relationship between IL-10 and alveolar macrophages, as this cell type is involved in inflammation in other virus infections of the lungs (*21, 22*). First, we examined the expression levels of various molecules associated with SARS-CoV-2 infection and found that the expression levels of *Ace2* and *Furin*, a protease essential for virus-binding to ACE2 (*23, 24*) and critical for SARS-CoV-2 entry, was significantly increased in alveolar macrophages skewed to M2c-type macrophages after IL-10 stimulation (**Fig. 2A**). In addition, the gene expression levels of the lectin receptor Siglec1 and the Fcgr1 Fc receptor, which were recently reported to be associated with SARS-CoV-2 infection and disease progression (*25, 26*), were also significantly elevated in IL-10-induced M2c-type alveolar macrophages (**Fig. 2B**). Furthermore, gene expression of the IL-6 receptor (Il6r), a key molecule in cytokine storms (*6*), was significantly increased in IL-10-induced M2c-type alveolar macrophages (**Fig. 2B**). However, the gene expression levels of CD147 (*27, 28*), Neuropilin-1 (Nrp1) (*29*), and Tmprss2 (*30*), which were reported to be important for virus entry, were unchanged in IL-10-induced M2c-type alveolar macrophages (**Fig. 2A**). These results suggested that IL-10 changed the phenotype of alveolar macrophages to M2c-type macrophages, which can be infected by SARS-CoV-2. Furthermore, ACE2 expression on M2c-type alveolar macrophages was inhibited by IL-10 receptor (IL-10R) blockade (**Fig. 2C**). A previous study reported that ACE2 expression was induced by interferon (IFN) in the airway epithelium (*31*), but this was not confirmed in alveolar macrophages, suggesting the manner of ACE2 induction is cell-specific (**Fig. 2C**). When IL-10 was administered intratracheally into mice *in vivo*, the expression of ACE2 on alveolar macrophages was increased compared with control mice (**Fig. 2D**). In contrast, experiments using epithelial and stromal cells showed no upregulation of ACE2 in response to IL-10 (**fig. S3**). In addition, IL-10-induced M2c-type alveolar macrophages exhibited proliferative and morphological changes (**fig. S4, A to D**). We performed an *in vitro* infection experiment to evaluate whether the ability of cells to be infected with SARS-CoV-2 increased under IL-10 stimulation. Importantly, SARS-CoV-2 rarely infected normal alveolar macrophages directly *in vitro* (**Fig. 2E**), whereas the SARS-CoV-2 virus was detected and proliferated in alveolar macrophages in *in vivo* infection experiments, indicating alveolar macrophages are changed by a factor released from the surrounding environment under the influence of SARS-CoV-2 infection (**fig. S4E**). Thus, we next performed infection experiments using IL-10-induced M2c-type macrophages and found that the expression of viral genomic RNA (N gene) increased in IL-10-treated cells (**Fig. 2E top**). Infecting virus RNA proliferated in M2c-type alveolar macrophages (**Fig. 2E bottom**), and fluorescent staining clearly showed that ACE2-expressing IL-10-induced M2c-type alveolar macrophages were infected with SARS-CoV-2 (**Fig. 2F**). These results suggested that IL-10, which is prevalent in patients with severe COVID-19, skewed alveolar macrophages towards an M2c-type macrophage that is susceptible to SARS-CoV-2, leading to increased inflammation within the alveoli.

**Fig. 2.**
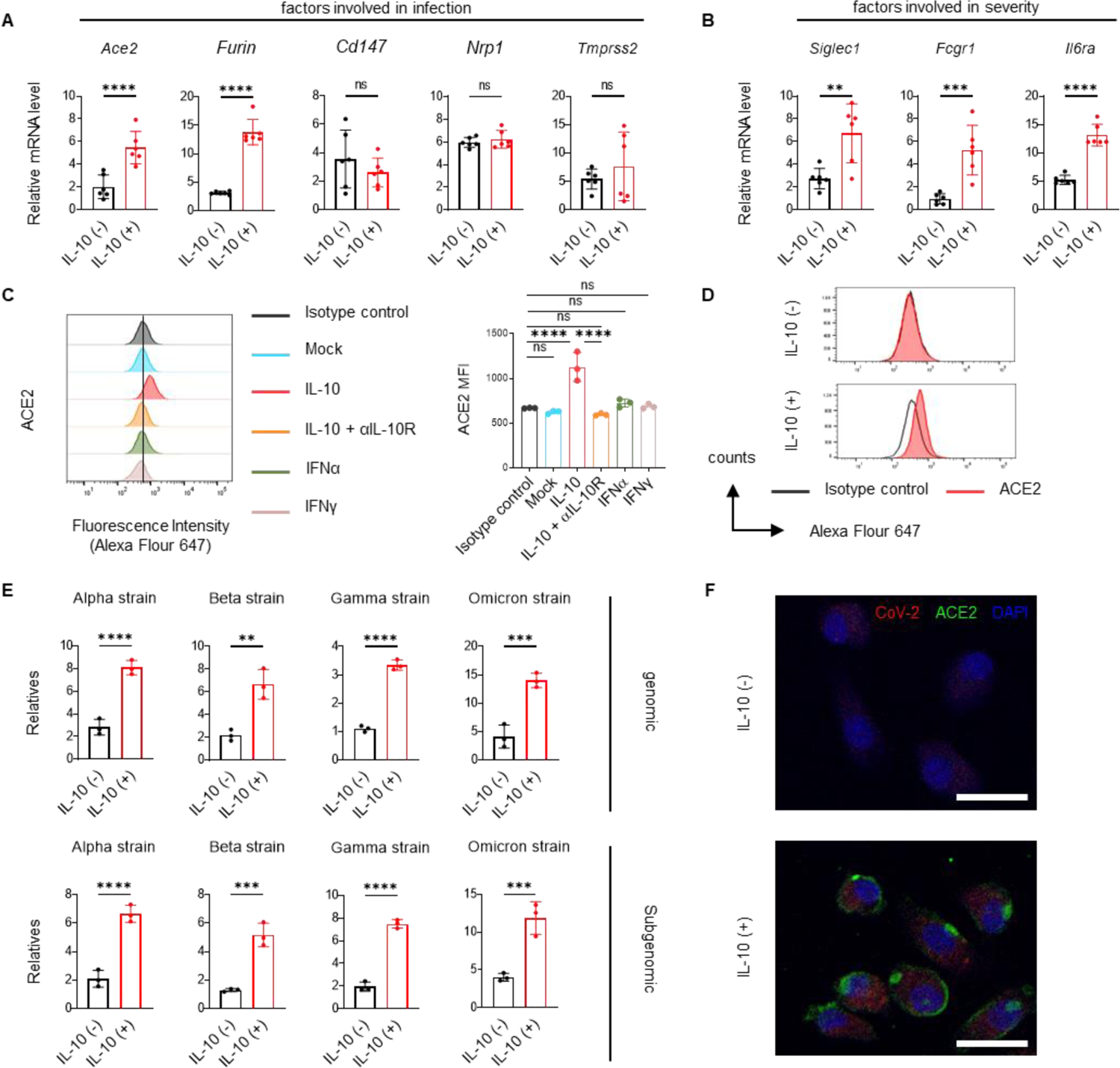
IL-10 promotes SARS-CoV-2 infection in alveolar macrophages. **(A and B)** mRNA expression of genes related to COVID-19 susceptibility **(A)** and severity **(B)** in murine alveolar macrophages stimulated by 40 ng/ml IL-10 for 24 h (n=6). **(C)** Flow cytometric analysis of ACE2 expression in murine alveolar macrophages stimulated by IL-10 (40 ng/ml) with and without IL-10R antibody (clone 1.3B1a, 10 µg/ml), IFN-α (40 ng/ml), and IFN-γ (40 ng/ml) for 72 h. **(D)** Flow cytometric analysis of ACE2 expression in alveolar macrophages isolated from IL-10-challenged mice at day 3 after intratracheal injection. **(E)** Quantitative PCR (qPCR) analysis of SARS-CoV-2 genomic and subgenomic (N) RNA in murine alveolar macrophages treated with IL-10 (40 ng/ml) 72 h after viral challenge (n=3). The error bars indicate mean ± standard deviation (SD). Statistical significance was determined by two-tailed, unpaired parametric *t*-tests **(A, B, and E)** or one-way ANOVA with Tukey multiple comparisons **(C)**. Data are representative of at least three independent experiments. **(F)** Immunofluorescence of infected alveolar macrophages with IL-10 (40 ng/ml) 72 h after viral challenge. Cells were stained for SARS-CoV-2 (Alexa Fluor 568, red), ACE2 (Alexa Fluor 488, green), and with DAPI (blue). Scale bars, 50 µm.

### Inhibition of M2c-skewing of alveolar macrophages induced by IL-10 attenuates the pathogenesis of COVID-19

To evaluate the relationship between M2c-type alveolar macrophages and the pathology of COVID-19, we developed an alveolar macrophage depletion model using the intratracheal administration of clodronate liposome (CLP) (**fig. S5, A and B**). On day 5 after COVID-19 infection, the expression levels of the inflammatory cytokine *Il6* and IFN-inducible *Cxcl10* and *Ccl5* significantly decreased in the lungs of the macrophage-depleted group (CLP group) compared with the control group injected with control liposomes (Control group) (**Fig. 3A**). Although there was no significant difference in the amount of viral RNA found in quantitative PCR (qPCR) analysis (**Fig. 3B**), we confirmed the virus was histologically localized in the central respiratory tract and that inflammation in the lung parenchyma was attenuated in the CLP group, suggesting that inflammation had not spread because the virus had not proliferated in the alveoli (**Fig. 3C**). In addition, to demonstrate that IL-10 promoted the exacerbation of alveolar-macrophage-mediated infection, we blocked IL-10R on the cell surface of alveolar macrophages. We used a continuous injection protocol based on flow cytometric analysis of IL-10R blocking (**fig. S5, C and D**) and found that the expression of inflammatory cytokines and IFN-inducible genes was significantly suppressed after IL-10R blocking in pulmonary tissues, like what was observed in the CLP group (**Fig. 3D**). In addition, inflammatory lesions in the lung parenchyma were reduced, and the virus accumulated in the central bronchus, although there was no significant difference in the quantities of viral RNA (**Fig. 3, E and F**). RNA-seq data from isotype control and IL-10R blocked groups after SARS-CoV-2 challenge showed that pathways associated with inflammatory response and IFN signaling were down-regulated in the IL-10R-blocked group (**Fig. 3G, fig. S5E, and table S4**). These results clearly showed that IL-10/IL-10R blockade in alveolar macrophages attenuated the pathology of COVID-19 infection. Our *in vitro* and *in vivo* studies on the relationship between IL-10 and alveolar macrophages revealed the mechanism by which macrophages acquire susceptibility to SARS-CoV-2 infection (**Fig. 3H**).

**Fig. 3.**
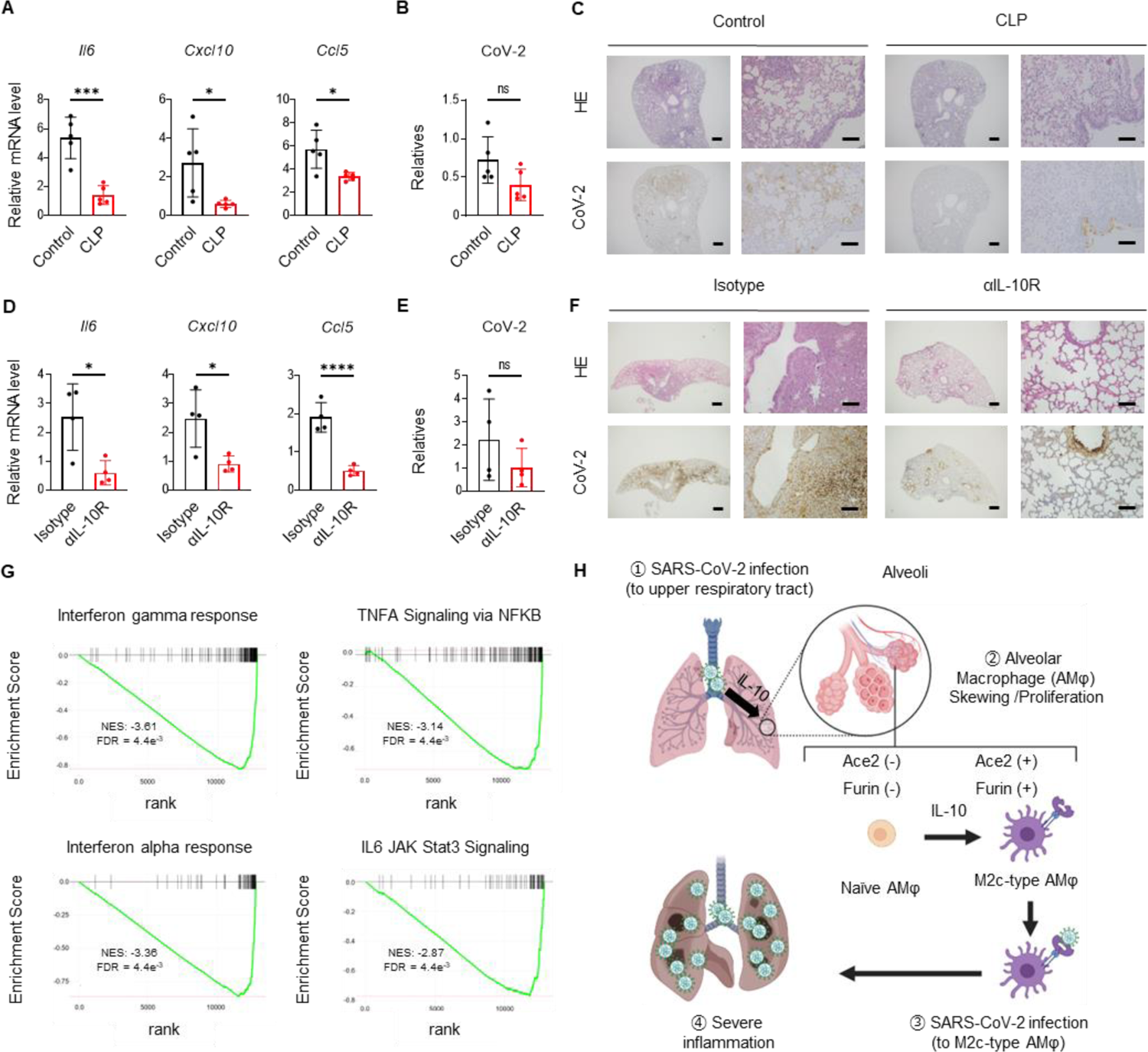
IL-10-activated alveolar macrophages trigger SARS-CoV-2 infection and exacerbate systemic inflammation in lungs. **(A to C)** Quantitative and histological analysis in a hamster alveolar macrophage depletion model between two groups: Control and CLP (n=5). **(A)** mRNA expression of inflammatory cytokines and chemokines. **(B)** Relative RNA levels of the SARS-CoV-2 genome. **(C)** HE (top) and immunohistochemistry of SARS-CoV-2 (bottom) staining of hamster lung specimens. Scale bars, 500 µm (left) and 100 µm (right) in both groups. **(D to F)** Quantitative and histological analysis in the IL-10R antibody administration model in hamsters between two groups: Isotype and anti-IL-10R (n=4). **(D)** mRNA expression of inflammatory cytokines and chemokines. **(E)** The relative RNA levels of the SARS-CoV-2 genome. **(F)** HE (top) and immunohistochemistry of SARS-CoV-2 (bottom) staining of hamster lung specimens. Scale bars, 500 µm (left) and 100 µm (right) in both groups. The error bars indicate mean ± SD. Statistical significance was determined by two-tailed, unpaired parametric *t*-tests **(A, B, D, and E)**. Data are representative of at least three independent experiments. **(G)** RNA-seq analysis of the IL-10R antibody administration model in hamsters (n=2). Enrichment plots of significantly enriched signaling pathways (top four) between the Isotype and anti-IL-10R groups are shown. Differences were considered significant when false discovery rate (FDR) < 0.05. **(H)** Graphical abstract of IL-10 signaling skewing alveolar macrophages towards the M2c phenotype, which leads to COVID-19 susceptibility and severity. Created with BioRender.com.

### Genetic analysis revealed novel mRNA transcripts generated by alternative splicing between *IFNAR2* and *IL10RB* genes were involved in human COVID-19 severity

It was suggested that IL-10 signaling in alveolar macrophages in the lungs is critical for COVID-19 aggravation. This prompted us to look for a genetic basis for the differences in IL-10 and IL-10R gene expression related to COVID-19 severity in humans. We investigated comprehensive gene expression data from many COVID-19 patient specimens. We evaluated the *IL10* and *IL10R* loci using genome-wide association study (GWAS) datasets obtained from the COVID-19 Host Genetics Initiative (HGI) (*32, 33*). GWAS are widely used to determine which genotypes in humans are likely to be associated with disease severity. First, we examined single-nucleotide polymorphisms (SNPs) around the *IL10* promoter region to confirm the association between patients with high IL-10 and COVID-19 severity. Although an SNP at the *IL10* promoter was reported to affect IL-10 expression *in vitro* (*34*), consistent with the expression quantitative trait loci (eQTL) effects for *IL10* on rs1800871 (**fig. S6A)**, no GWAS signal associated with COVID-19 severity was found in the same region (**fig. S6B**). Next, we focused on IL-10R, which amplifies IL-10 signals and consists of IL-10RA and IL-10RB subunits. Regarding IL-10RA, GWAS signals in this region were not involved in COVID-19 severity (**fig. S6C**). Next, we examined SNPs at the region around the *IL10RB* locus on chromosome 21. GWAS demonstrated the *IFNAR2* locus located near *IL10RB* was significantly associated with COVID-19 severity, consistent with recent reports (*35–37*) (**fig. S6D**). We evaluated the cell-specific eQTL effects of *IL10RB* expression on monocytes/macrophages according to rs13050728 (Variant ID; chr21:33242905:T:C), which had the most significant association with COVID-19 severity on the locus (*P* = 4.07e^-21^, T = risk allele) in the COVID-19 HGI A2 datasets (**Fig. 4A**), whose QTL data were obtained from two cohorts (the DICE and EvoImmunoPop project) (*38–41*). There was no significant eQTL effects on *IL10RB* expression in these datasets, indicating that this variant does not affect the IL-10RB expression pattern in monocytes/macrophages (**Fig. 4B**). In contrast, we found a unique splicing QTL (sQTL) effect on rs13050728 at the splice junction between *IFNAR2* and *IL10RB* (chr21: 33252830: 33268394: clu_34140_+) in monocyte/macrophage datasets (**Fig. 4C**). Thus, we performed co-localization analysis between this sQTL and GWAS for COVID-19 severity. Interestingly, the sQTL effect according to rs13050728 in monocyte/macrophage datasets was significantly co-localized with the GWAS results (*PP-H4* = 0.96), indicating that patients harboring the risk (T) allele of rs13050728 had a higher proportion of the novel splicing isoforms and were more likely to develop severe disease compared with those with the nonrisk (C) allele (**Fig. 4D**). To investigate which transcripts were expressed under the influence of this sQTL effect, we performed long-read capture sequencing using the monocyte/macrophage transcriptome. We identified two novel transcripts in addition to the normal *IFNAR2* and *IL10RB* mRNA transcripts (**Fig. 4A and table S5**). These novel transcripts were fused with parts of *IFNAR2* and *IL10RB* as a result of the unique splicing and contained the same coding sequence (CDS) (**fig S6E and table S5**). Thus, we evaluated the expression levels of the novel transcripts in human peripheral blood mononuclear cell (PBMC)-derived monocytes/macrophages. Intriguingly, the novel transcripts showed a significant increase in the T/T genotype compared with the C/C and C/T genotypes (**Fig. 4E**). However, *IL10RB* expression was lower in the T/T genotype compared with the C/C genotype, and *IFNAR2* expression levels were not significantly different in any genotype (**Fig. 4E**). These results clearly indicated that the risk (T) allele of rs13050728 increased the amount of the novel transcripts in monocytes/macrophages, and the expression levels of the novel transcripts was strongly correlated with COVID-19 infectivity and severity in humans.

**Fig. 4.**
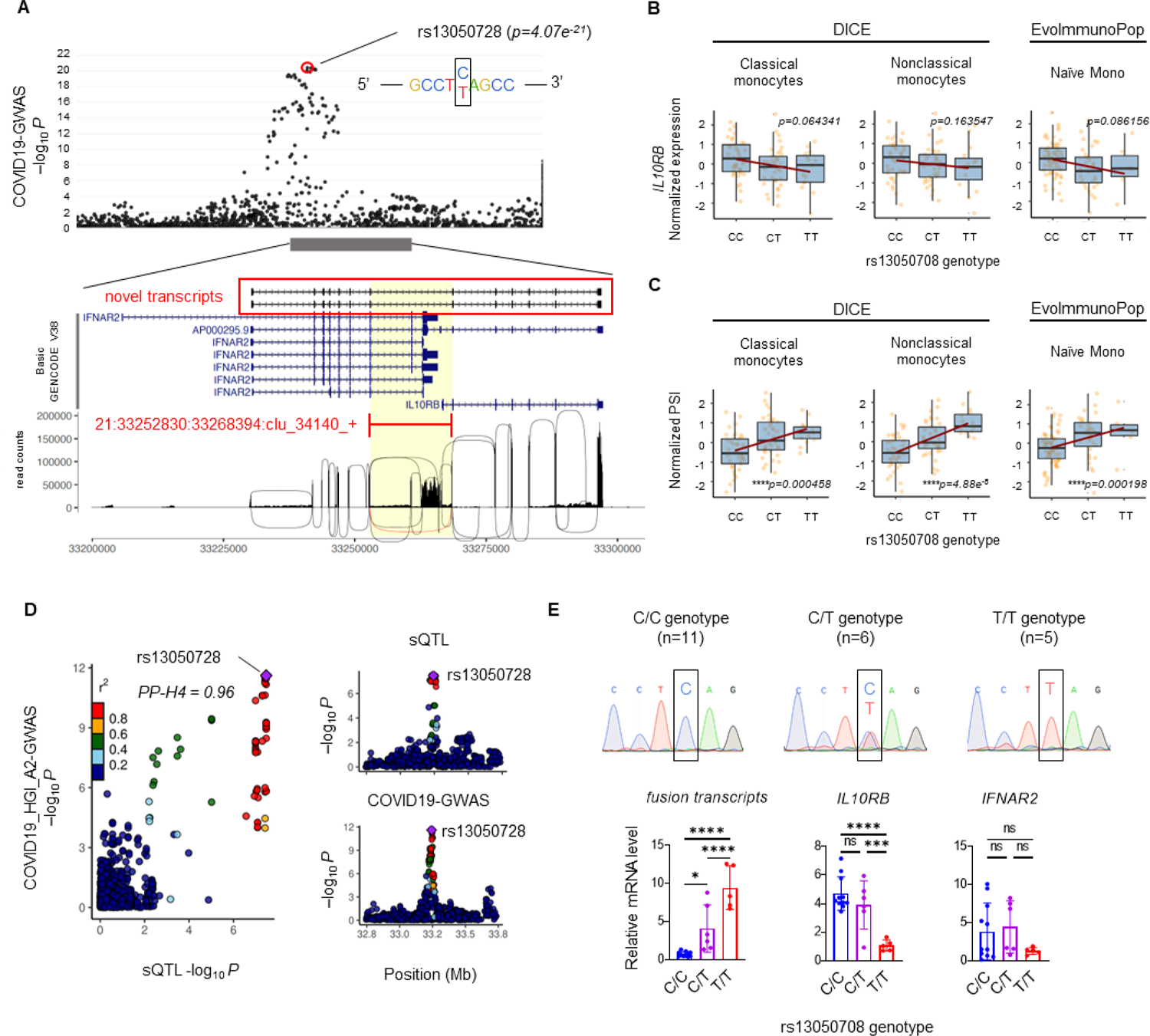
*IFNAR2-IL10RB* fusion transcripts are highly expressed in COVID-19 patients with COVID-19 severity risk-allele at rs13050728. Genetic analysis of human monocytes/macrophages based on rs13050728 genotypes, which were significantly associated with COVID-19 severity in COVID-19 HGI Release 5 GWAS (A2 datasets: Critically ill COVID-19 patients vs population controls). The rs13050728 T allele is a risk allele for COVID-19 severity. **(A)** Diagram showing the results of the genetic analysis of the *IFNAR2*-*IL10RB* locus. Manhattan plot shows GWAS results from COVID-19 HGI Release 5 A2 datasets. SNPs in the *IFNAR2-IL10RB* locus are shown as dots. The picture below shows the exon containing the novel transcripts (red square) and other isoforms annotated in Basic GENECODE V38. The splicing pattern and read counts in the same locus are shown. A splicing pattern affected by rs13050728 variants (21:33252830:33268394:clu_34140_+) is shown as a red line. **(B and C)** QTL analysis of monocyte/macrophage datasets based on the rs13050728 genotype. SNP array and RNA-seq data obtained from the DICE and EvoImmunoPop project were used. Data were analyzed using a linear regression model with MatrixQTL (*66*). **(B)** Box plot shows the eQTL effect on *IL10RB* expression. **(C)** Box plot shows the sQTL effect on the intron excision of 21:33252830:33268394:clu_34140_+. Normalized gene expression levels and percent spliced in (PSI) were plotted based on the genotype of rs13050728, respectively. Each box reflects the interquartile range (IQR), and the upper and bottom whiskers represent the maximum and minimum values within 1.5 times IQR from the hinge, respectively. **(D)** Colocalization analysis of the sQTL effects on the intron excision of 21:33252830:33268394:clu_34140_+ with a GWAS signal for COVID-19 in the same region. SNPs on the *IFNAR2-IL10RB* locus are represented as dots colored based on linkage disequilibrium (r^2^). rs13050728 is marked with a purple dot. PP-H4 colocalization is also indicated. The DICE nonclassical monocyte datasets were used. **(E)** Relative expression of the *IFNAR2*-*IL10RB* fusion transcripts, *IL10RB* and *IFNAR2* in human PBMC-derived monocytes/macrophages according to the rs13050728 genotype (n=22). The error bars indicate mean ± SD. Statistical significance was determined by one-way ANOVA with Tukey multiple comparisons.

### *IFNAR2/IL10RB*-fused transcripts encode unique “hybrid” receptor

Based on the common CDS of the novel transcripts (**fig. S6E**), we examined its corresponding amino acid sequence and found it contained the extracellular domains of IFNAR2 and IL-10RB, and the transmembrane and intracellular domains of IL-10RB (**Fig. 5, A and B**). To investigate whether the transcripts were successfully translated and translocated to the cell surface, the full-length CDS was overexpressed in HEK293 cells. As a result, we confirmed that the coding protein was translated and translocated to the cell surface (**fig. S6, F and G**). Next, we investigated whether complexes of hybrid receptors and other receptors could bind to their corresponding ligands. The wild-type IL-10RB and IFNAR2 formed complexes with IL-10RA and IFNAR1, respectively, for ligand recognition. The IL-10RB/IL-10RA complex forms a dimer that interacts with dimeric IL-10, whereas the IFNAR2/IFNAR1 complex forms a monomer that interacts with monomeric IFN-α (*42, 43*). Thus, we modelled a complex of the hybrid receptor, which contains IL-10RB and IFNAR2, with IL-10RA as a dimer or IFNAR1 as a monomer using AlphaFold v2.2.2. From a structural point of view, the Hybrid/IL-10RA successfully formed a complex that could activate downstream IL-10 signaling pathways (**Fig. 5C**). In contrast, the Hybrid/IFNAR1 was predicted not to be activated by IL-10 signaling because each intracellular domain was too far apart (**Fig. 5D**). Next, we evaluated the binding affinities of the Hybrid complex with ligands, including IL-10 and IFN-α, using the DockQ (pDockQ) score program (detailed in **Materials and Methods**). The binding affinities of complexes containing the Hybrid/IL-10RA receptor with the ligand IL-10 were predicted to be similar to those of wild-type IL-10RB/IL-10RA, whereas most complex models involving IFN-γ, a negative control ligand, had quite low pDockQ scores, suggesting that the Hybrid/IL-10RA and Hybrid/IFNAR2 did not recognize IFN-γ (**Fig. 5E**). In addition, the pDockQ score for Hybrid/IFNAR2 and IFN-α (a known IFNAR2 ligand) was also high, indicating that the Hybrid receptors could bind to IFN-α but were unable to activate downstream signaling pathways (**Fig. 5, D and E**). Since this novel receptor was suggested to recognize two molecules, we termed this “COVID-19 infectivity enhancing dual receptor” (CiDRE).

**Fig. 5.**
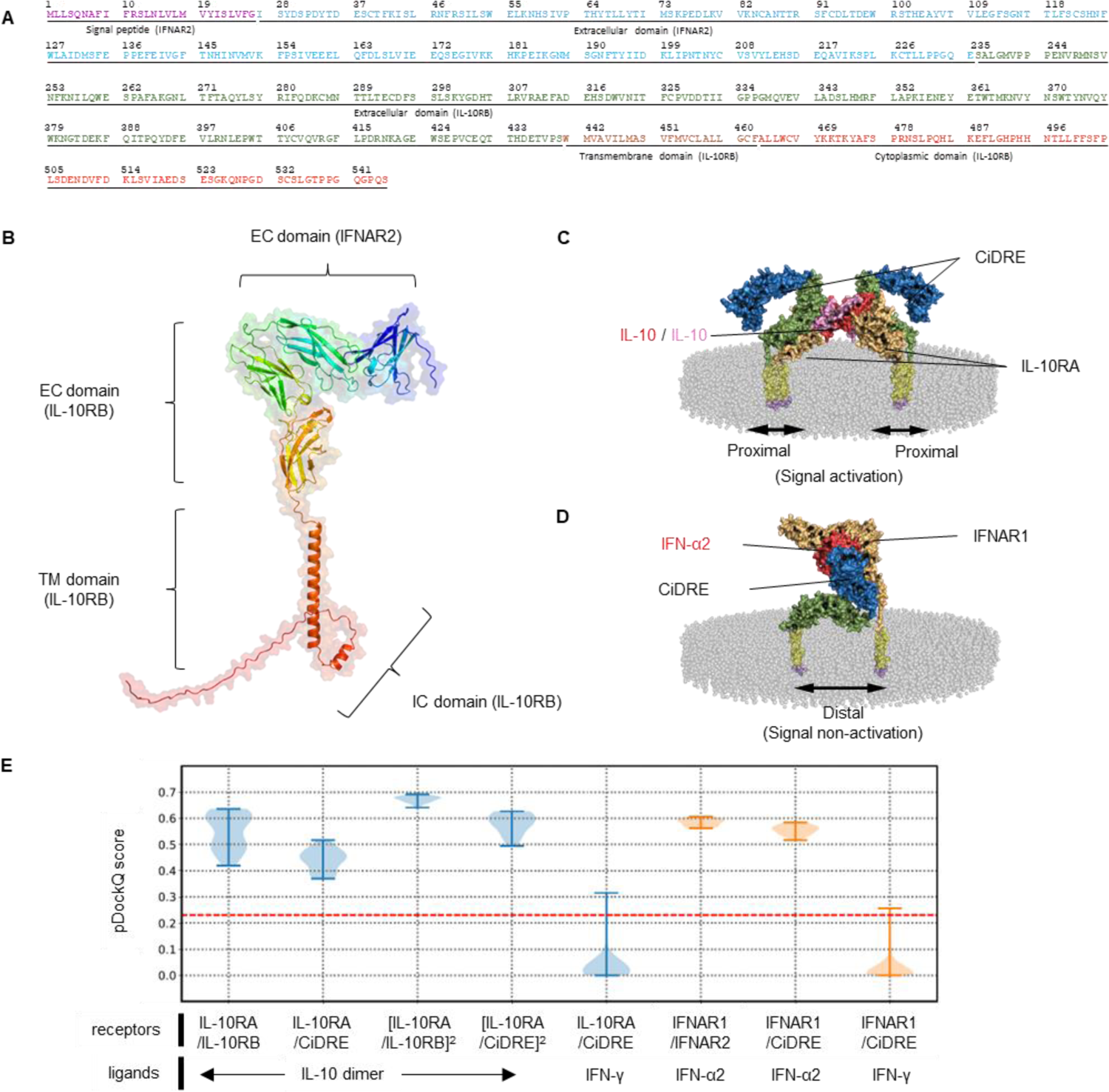
A computational structure model of the hybrid receptor. **(A)** Amino acid sequence of the hybrid receptor predicted from the CDS of the *IFNAR2*-*IL10RB* fusion transcripts. The functional domains predicted from the wild-type IFNAR2 and IL-10RB subunits are shown as purple (signal peptide), blue (extracellular domain of IFNAR2), green (extracellular domain of IL-10RB), orange (transmembrane domain of IL-10RB) and red (cytoplasmic domain of IL-10RB). **(B)** Structural modelling of the IFNAR2-IL-10RB hybrid receptor. The hybrid receptor model covers residues 27–545. The signal peptide at the N-terminus was excluded from modelling. The chain colors from the N-terminus (blue) to the C-terminus (red) vary with a rainbow spectrum. **(C)** Structural modelling of the IL-10RA-Hybrid receptor in a complex with an IL-10 dimer embedded in a lipid bilayer membrane. The extracellular domains of IFNAR2 and IL-10RB within the hybrid receptor are shown as blue and green, respectively. Two IL-10RA receptors are shown in brown. Two dimerized IL-10 chains are shown as pink and red. Transmembrane domains are yellow, and short fragments of cytoplasmic domains are purple. The cell membrane is represented as translucent spheres in grey. **(D)** Structural modelling of the IFNAR1-Hybrid receptor in complex with IFN-α2 embedded into a lipid bilayer structure. The extracellular domains of IFNAR2 and IL-10RB are shown as blue and green. The IFNAR2 receptor is brown and the IFN-α2 molecule is red. Transmembrane domains are yellow, and the short fragments of cytoplasmic domains are purple. **(E)** The binding affinity of different receptor/ligand complexes measured by pDockQ scores. pDockQ score threshold (0.23) is plotted as a dashed red line, above which a complex model was categorized as acceptable and the higher the better.

### CiDRE is involved in COVID-19 severity as a dual receptor for IL-10 and type I IFN signaling

To investigate the biological function of human CiDRE in alveolar macrophages, murine alveolar macrophages transduced with human CiDRE-expressing lentiviral vectors were stimulated with cytokines, including human IL-10 (hIL-10), and the phosphorylation of STAT3 was examined, since STAT3 is phosphorylated in response to IL-10 (*44*). STAT3 phosphorylation was strongly amplified in hIL10-stimulated CiDRE-expressing macrophages compared with controls, consistent with the model of hIL-10 binding to hIL-10R and mouse IL-10R (mIL-10R) (**Fig. 6A**). In contrast, the phosphorylation of STAT3 and its amplification were not found in response to human IFN-α (hIFN-α) or IFN-γ (hIFN-γ). In addition, augmented phosphorylation via CiDRE in response to hIL-10 was suppressed by the addition of an anti-human IL-10RB antibody (clone 90220) that recognizes hIL-10RB but not mIL-10RB (**Fig. 6B**). Moreover, the expression levels of *Ace2* was significantly augmented by IL-10 in CiDRE-expressing alveolar macrophages (**Fig. 6C**). Although type I IFN did not activate IL-10 signaling in CiDRE-expressing macrophages, IFN-α was strictly associated with CiDRE (**Fig. 6, A and D**). Taken together, the evidence suggests CiDRE and IL-10R bind strongly to IL-10 to synergistically activate M2c-type macrophages, which can be infected by SARS-CoV-2, whereas CiDRE functions as a decoy receptor that sequesters type I IFNs, which have important roles in antiviral responses.

**Fig. 6.**
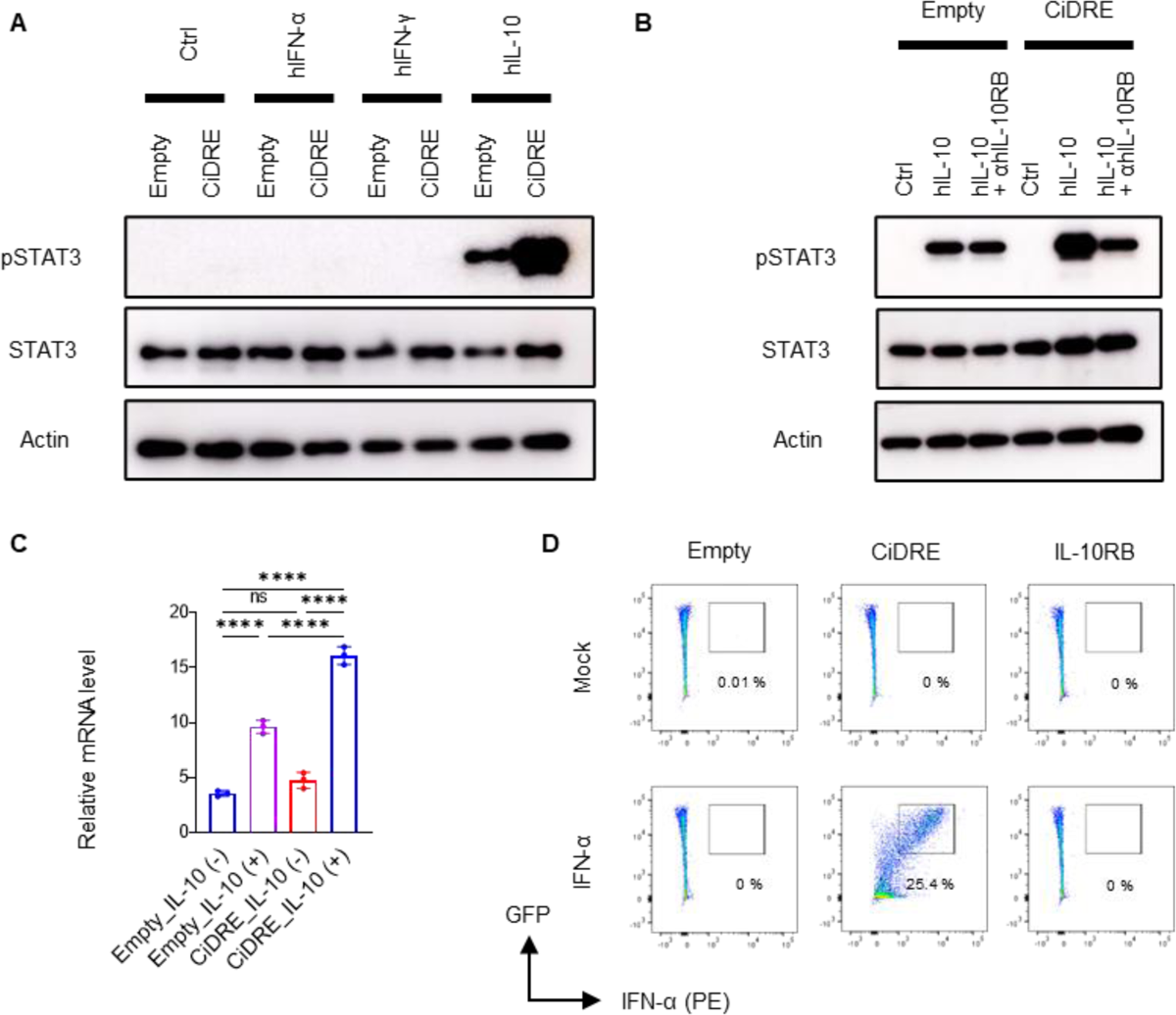
CiDRE is a dual-receptor related to COVID-19 exacerbation. **(A and B)** Western blot (WB) analysis of CiDRE-expressed murine alveolar macrophages. As a control, alveolar macrophages transduced with empty lentiviral vectors were used. **(A)** WB analysis of CiDRE-expressing murine alveolar macrophages treated with 40 ng/ml of hIFN-α, hIFN-γ, or hIL-10. **(B)** WB analysis of CiDRE-expressing murine alveolar macrophages treated with 40 ng/ml hIL-10 and 1 µg/ml anti-human IL-10RB antibody (clone 90220), which reacts only with human CiDRE but not native murine IL-10RB. **(C)** mRNA expression of ACE2 in CiDRE-expressing alveolar macrophages stimulated by 40 ng/ml hIL-10 for 24 h (n=3). The error bars indicate mean ± SD. Statistical significance was determined by one-way ANOVA with Tukey multiple comparisons. **(D)** Flow cytometric analysis of CiDRE-expressing murine alveolar macrophages. CiDRE-expressing cells were stimulated by 40 ng/ml hIFN-α followed by incubation with anti-human IFN-α antibody (PE conjugated). IFN-α-binding activity was evaluated by the fluorescent intensity of PE in CiDRE-expressing cells, which were green fluorescent protein (GFP) positive. Data are representative of at least three independent experiments.

Taken together, our findings confirmed an antibody-independent pathway of SARS-CoV-2 infection in M2c-type alveolar macrophages and clarified the mechanism of COVID-19 severity mediated by the IL-10 pathway. In addition, analysis using human samples demonstrated a genetic difference in IL-10 reactivity in alveolar macrophages that was affected by the expression levels of the novel receptor, CiDRE.

## Discussion

We found that IL-10 signal amplification mediated by the native IL-10RB and CiDRE, followed by M2c-skewing of alveolar macrophages, is critical for disease progression in COVID-19. This finding may have important implications for predicting the severity of COVID-19, prioritizing vaccination programs, and developing new preventive drugs such as anti-IL10R inhalers. Recent studies reported the involvement of monocytes/macrophages in COVID-19 (*15, 16, 45*). Monocytes/macrophages were reported to be diverse, and resident macrophages and bone marrow-derived macrophages are thought to contribute to the pathogenesis of lung disease (*46, 47*). Spatiotemporal analysis of these myeloid cells is very important, and the relationship between IL-10 and alveolar macrophages in this study suggested they are responsible for the transition from the early stages of infection to a progressive state and, therefore, might determine the fate of patients with SARS-CoV-2. We confirmed that the expressions of IL-10RB and CiDRE are mutually exclusive, indicating that the upregulation of CiDRE influences the total IL-10 signal intensity because IL-10 was reported to have a low binding affinity for IL-10RB (*42*). Therefore, small changes in the ligand-receptor relationship can have a large impact on the downstream signaling pathways (*42*). In addition, CiDRE is expected to act as a decoy receptor for type I IFNs, thereby delaying viral clearance and promoting the progression of COVID-19. Interestingly, compared with SARS and Middle East respiratory syndrome, COVID-19 was suggested to significantly increase Th2 cytokines, including IL-10 (*48*). CiDRE might be specific to SARS-CoV-2 infection; however, it might also be involved in IL-10-related diseases such as Crohn’s disease, ulcerative colitis, and psoriasis (*49*). Furthermore, CiDRE might have dominant functions in other immune cells that have low expression levels of IL-10RB, such as lymphocytes. CiDRE represents a paradigm shift in our understanding of the relationship between type I IFN and IL-10 signaling and their roles in the development of disease.

## Materials and Methods

### Animals

BALB/c mice and Syrian hamsters were purchased from Japan SLC (Shizuoka, Japan). Male mice between 6 and 8 weeks of age and male hamsters between 3 and 4 weeks of age were used. All animals were housed in individually ventilated cages with free access to food and water in a temperature and light-regulated room in a specific pathogen free (SPF) facility. All animal experiments without SARS-CoV-2 were performed at the Tokyo Medical and Dental University. The study protocols were approved by the ethical committees of Tokyo Medical and Dental University (A2021-275C and A2021-294A). All animal experiments with SARS-CoV-2 were performed in Animal Biosafety Level 3 (ABSL3) facilities at the Research Institute for Microbial Diseases, Osaka University. The study protocols were approved by the Institutional Committee of Laboratory Animal Experimentation of the Research Institute for Microbial Diseases, Osaka University (R02-08-0). All efforts were made during the study to minimize animal suffering and to reduce the number of animals used in the experiments.

### Cells and Viruses

HEK293 (ATCC, Manassas, VA, USA), HEK293T (ATCC), and A549 (ATCC) cells were cultured in Dulbecco’s Modified Eagle Medium (DMEM) supplemented with 10% heat-inactivated foetal bovine serum and 1% penicillin/streptomycin (complete DMEM). Roswell Park Memorial Institute (RPMI) 1640 medium was used for THP-1 (ATCC) cells instead of DMEM. MLE12 cells and primary lung fibroblasts were cultured as described previously(*50, 51*). Vero E6/TMPRSS2 cells were provided by the National Institutes of Biomedical Innovation, Health and Nutrition (Japan) and maintained at 37°C with 5% CO_2_ in complete DMEM. SARS-CoV-2 strains, hCoV-19/Japan/QHN002/2021 (Alpha), hCoV-19/Japan/TY8-612/2021 (Beta), hCoV-19/Japan/TY7-503/2021 (Gamma), hCoV-19/Japan/TY11-927/2021 (Delta), and 2019-nCoV/Japan/TY38-873/2021 (Omicron) were isolated at the National Institute of Infectious Diseases (Japan) and provided for this research. SARS-CoV-2 was propagated in VeroE6/TMPRSS2 cells. The virus stock was generated from the supernatant of VeroE6/TMPRSS2 cells infected with SARS-CoV-2 at a multiplicity of infection (MOI) of 0.1 and harvested 2 days after infection. The viral titer was determined by plaque assay.

### Human clinical samples

This study was in part based on clinical materials and information from the BioBank at the Bioresource Research Center, Tokyo Medical and Dental University (BRC2021-001). Human serum samples for ELISA were obtained from 122 hospitalized patients with COVID-19 at the Tokyo Medical and Dental University. All subjects signed a consent form approved by the ethical committees of Tokyo Medical and Dental University. Ethical approval for the human study protocols was obtained from the ethical committees of Tokyo Medical and Dental University (G2020-034).

### Gene cloning

The coding sequence (CDS) of the CiDRE transcripts was amplified by PCR using KOD FX Neo (TOYOBO, Osaka, Japan) with THP-1 complementary DNA as a template and the following primer pairs: Forward: 5′-ATGCTTTTGAGCCAGAATGCCTTC-3′ and Reverse: 5′-CTAGCTTTGGGGCCCCTGCCCA-3′. The PCR reaction was performed with the following settings: initial denaturation step of 4 min at 96°C, amplification of 40 cycles of 30 s denaturation at 96°C, followed by annealing for 30 s at 70°C and the last extension step of 7 min at 72°C. Blunt-end amplified CDS products were cloned into a pCR-Blunt II-TOPO vector using a Zero Blunt™ TOPO™ PCR Cloning Kit (Thermo Fisher Scientific, Waltham, MA, USA) in accordance with the manufacturer’s protocol. The cloned vectors were sent to Azenta Japan Corp. (Tokyo, Japan) and sequenced with two universal primers (M13 Forward and M13 Reverse) to confirm the CiDRE-CDS total sequence. Then, a PCR reaction was performed in the same manner using a vector including the correct CiDRE-CDS sequence with the following primer pairs: Forward: 5′-GGATCCATGCTTTTGAGCCAGAATGCC-3′ and Reverse: 5′-CTCGAGCTAGCTTTGGGGCCCCTG-3′, which were designed to have *Bam*HI and *Xho*I restriction sites on the 5′ end of the former primers, respectively. PCR products were cloned into a subcloning vector and then the CiDRE-CDS insert sequence was transferred to a FUGW-IRES-GFP (FUIGW) vector (*52*) after enzymatic restriction by *Bam*HI (TOYOBO) and *Xho*I (TOYOBO), followed by ligation using a DNA Ligation Kit Ver. 2.1 (Takara, Shiga, Japan). In this experiment, competent high DH5α (TOYOBO) competent cells were used for plasmid amplification.

### Transfection

HEK293 cells were transfected with FUIGW-NanoLuc (empty control), FUIGW-CiDRE, or FUIGW-IL-10RB using Lipofectamine 2000 (Thermo Fisher Scientific) following the manufacturer’s protocols. Then, 48 h after transfection, cells were incubated with 40 ng/ml human IL-10 (BioLegend, San Diego, CA, USA) for 2 h at 4°C and cells were collected for flow cytometric analysis.

### Lentiviral transduction

Lentiviruses were produced in HEK293T cells by the co-transfection of FUIGW-NanoLuc or FUIGW-CiDRE with two lentiviral helper plasmids, pCMV-dR8.2 (*53*) and pCAG-VSV-G (*53, 54*). Culture supernatants containing lentivirus were collected 24 and 48 h after transfection and concentrated by a Lenti-X concentrator (Takara) followed by passing through a 0.45-μm PES filter. Lentiviral transduction of murine alveolar macrophages was performed for 7 days in the presence of 5 μg/cm^2^ RetroNectin (Takara). Stable gene expression was confirmed by GFP signals using a BZ-X710 (KEYENCE, Osaka, Japan). For WB analysis, successfully transduced cells were collected after 10 min incubation with human IFN-α (BioLegend), IFN-γ (BioLegend), or IL-10 with or without anti-IL-10RB antibody (clone 90220, R&D Systems, Minneapolis, MN, USA) for 30 min. For qPCR analysis, cells were collected 24 h after incubation with 40 ng/ml IL-10 following the protocol of the High Pure RNA Isolation Kit (Roche, Basel, Switzerland).

### Intratracheal injection of IL-10

BALB/c mice were anaesthetized by the intraperitoneal administration with 10 µl/g body weight of a medetomidine-midazolam-butorphanol tartrate mixture (0.75 µg/ml medetomidine, 4 µg/ml midazolam, and 5 µg/ml butorphanol tartrate), and then the intratracheal administration of murine IL-10 (10 µg/animal in 50 µl phosphate buffered saline (PBS)) was performed. The intraperitoneal administration of atipamezole was performed for the recovery from anesthesia. Bronchoalveolar lavage (BAL) and lungs were collected at 3 days after instillation.

### BAL

To isolate alveolar macrophages, animals were euthanized by CO_2_ narcosis. Then, the lungs were inflated with 1-ml (mice) or 2.5-ml (hamsters) aliquots of PBS three times. After centrifugation and red blood cell lysis, the collected cells were cultured at 37°C with 5% CO_2_ in complete RPMI for 4 h. Then, the attached cells (typically >95% alveolar macrophages) were stimulated with 40 ng/ml murine IL-10, IFN-α, or IFN-γ (all from BioLegend). For IL-10R blocking, cells were treated with 10 µg/ml anti-IL-10R antibody (clone 1.3B1a, BioLegend) for 30 min prior to IL-10 stimulation. Then, samples were collected 72 h after incubation for flow cytometric analysis. For IL-10 intratracheal injection experiments, collected BAL samples were directly used for subsequent experiments. For SARS-CoV-2 challenge, isolated alveolar macrophages were infected with each strain of SARS-CoV-2 (Alpha, Beta, Gamma, and Omicron) at an MOI of 10 24 h after incubation with or without IL-10. Cells were harvested at 72 h after viral challenge and lysed for RNA extraction or fixed in 10% neutral-buffered formalin (NBF) for immunofluorescence staining.

### Preparation of primary lung cells

Murine lungs were dissociated with 1 mg/ml type I collagenase (Sigma, St. Louis, MO, USA) in the presence of DNase I (Roche) for 30 min at 37°C. The cells were then centrifuged at 300 × g for 10 min and treated with RBC Lysing buffer for red blood cell lysis for 2 min at room temperature. The collected samples were suspended in FACS buffer (0.5% bovine serum albumin (BSA) and 2 mM EDTA in PBS) for flow cytometric analysis.

### Syrian hamster COVID-19 model

Syrian hamsters were anaesthetized with isoflurane and challenged with 1.0 × 10^6^ plaque-forming unit SARS-CoV-2 (Delta strain) via the intranasal route. For alveolar macrophage depletion experiments, hamsters were anaesthetized with the anesthetic tartrate mixture as above, and then the intratracheal administration of clodronate liposome 100 (Katayama Chemical, Osaka, Japan) or control liposomes (Katayama Chemical) was performed 7 days prior to infection. For IL-10 blocking experiments, daily intratracheal injection with 400 µg/body of anti-IL-10R antibody (clone 1.3B1a, BioLegend) or isotype control (BioLegend) under anesthesia with the tartrate mixture was started 2 days post-infection until the day of analysis. Atipamezole was used for hamster recovery as described above. In both experiments, hamsters were euthanized by CO_2_ narcosis and lungs were collected 5 days post-infection for subsequent experiments.

### quantitative PCR

Total RNA was extracted using a High Pure RNA isolation kit (Roche) or TRIzol (Thermo Fisher Scientific), in accordance with the manufacturer’s instructions. If necessary, RNA was concentrated using a Monarch RNA Cleanup Kit (New England Biolabs, Ipswich, MA, USA). After reverse transcription with the ReverTra Ace qPCR RT Master Mix (TOYOBO), qPCR was performed using THUNDERBIRD SYBR qPCR Mix (TOYOBO) on a LightCycler 480 instrument (Roche). Relative values of target gene expression were calculated using the standard curve method normalized to β-actin. qPCR primer sequences are listed in **table. S6**.

### Flow cytometry

After Fc-blocking using FcX reagents (BioLegend), cells were incubated with primary antibodies in FACS buffer for 30 min at 4°C and if necessary, incubated with secondary antibodies for 20 min at 4°C. Lung epithelial cells were defined as CD45^−^CD31^−^EpCAM^+^ cells. Lung endothelial cells were defined as CD45^−^CD31^+^EpCAM1^−^ cells. Lung fibroblasts were defined as lung CD45^−^CD31^−^EpCAM1^−^CD140a^+^ cells. For the assessment of ACE2 expression, goat anti-ACE2 antibody (R&D Systems) and Alexa Flour 647-conjugated anti-goat IgG antibody (Abcam, Cambridge, UK) were used as primary and secondary antibodies, respectively. Data were acquired on a flow cytometer (FACS CantoII; BD Bioscience, Franklin Lakes, NJ, USA) and analyzed using FlowJo software (Tree Star, Inc., Ashland, OR, USA).

### ELISA

IL-10, IL-6, IFN-α, IFN-γ, IFN-λ, IL-1α, IL-1β, TNF-α, IL-2, IL-4, IL-5, IL-13, IL-33, MCP-1, HMGB1, TGF-β, NGAL, and MRP-8/14 in human blood serum were measured by ELISA kits (BioLegend, R&D Systems, or Shino-Test Corp., Tokyo, Japan) following the manufacturer’s protocols.

### Western blotting

Separation of protein samples, which were quantified using a Pierce BCA Protein Assay Kit (Thermo Fisher Scientific), was performed by SDS-PAGE. After transfer to polyvinylidene difluoride (PVDF) membranes, the membranes were incubated with the appropriate primary antibodies overnight. Then, membranes were incubated with HRP-conjugated secondary antibodies, and reacted with HRP substrates (Merck Millipore, Darmstadt, Germany) for enhanced chemiluminescence (ECL) detection using Amersham Imager 680 (Cytiva, Marlborough, MA, USA). β-Actin was selected as a loading control.

### Histopathology

Lung tissues were fixed in 10% NBF for 24–48 h and embedded in paraffin. Sections were cut at 4-µm thickness using a microtome and mounted on glass slides. After deparaffinization and dehydration, the slides were used for subsequent HE staining and other staining procedures. To evaluate fibrotic lesions, sections were stained with azan. Terminal deoxynucleotidyl transferase-mediated dUTP nick end labeling (TUNEL) staining was performed using an Apop Tag Plus Peroxidase In Situ Apoptosis Kit (Merck Millipore) following the manufacturer’s protocols. Histopathological evaluation was performed using a BZ-X710 microscope.

### Immunochemistry and Immunofluorescence

For immunochemistry and immunofluorescence, heat-activated antigen retrieval of deparaffinized/dehydrated sections was performed using Target Retrieval Solution (Dako, Glostrup, Denmark). After the blocking of endogenous peroxidases using Peroxidase-Blocking Solution (Dako), the sections were incubated overnight at 4°C with anti-ACE2 (1:100), mouse anti-SARS-CoV-2 (1:1000), and rabbit anti-LC3B (1:1000) antibodies in Antibody Diluent (Dako) as primary antibodies. If necessary, the M.O.M. Fluorescein Kit (Vector Laboratories, Newark, CA, USA) was used for endogenous mouse IgG blocking. For immunochemistry, the Envision+ System-HRP Labeled Polymer (Dako) was used as a secondary antibody. Counter staining was performed by hematoxylin followed by DAB staining. Then, histopathological evaluation was performed using a BZ-X710 microscope. For immunofluorescence, the appropriate fluorescent labelled antibodies were used as secondary antibodies with DAPI (Bio-Rad, Watford, England). The sections were incubated for 60 min at room temperature in the dark. Then, the histopathological evaluation was performed using an LSM 780 microscope (Carl Zeiss, Oberkochen, Germany).

### Electron microscopy

The specimens were fixed in 2.5% glutaraldehyde in 0.1 M phosphate-buffer (PB) for 2 h. The specimens were washed overnight at 4°C in 0.1 M PB and post-fixed with 1% osmium tetroxide buffered with 0.1 M PB for 2 h. For scanning electron microscopy, the specimens were dehydrated in a graded series of ethanol and dried in a critical point drying apparatus (JCPD-5; JEOL, Tokyo, Japan) with liquid CO_2_. The specimens were spatter-coated with platinum and examined by scanning electron microscope (JSM-7900F; JEOL). For transmission electron microscopy, the specimens were dehydrated in a graded series of ethanol and embedded in Epon 812 (TAAB Laboratories Equipment, Aldermaston, England). Ultrathin sections (70 nm thickness) were collected on coppergrids, double-stained with uranyl acetate and lead citrate and then examined by transmission electron microscope (JEM-1400Flash; JEOL). Experiments in electron microscopy were performed at the Research Core of Tokyo Medical and Dental University.

### Human PBMC and Genotyping

Human PBMCs were purchased from Precision for Medicine, Inc. (Cat# 551-37651 and 555-41341; Frederick, MD, USA) and cultured for 7 days with 20 ng/ml recombinant M-CSF (R&D Systems) to allow them to differentiate to macrophages. The macrophages were used for the qPCR analysis of CiDRE transcripts and genotyping at rs13050728. For genotyping at rs13050728, PCR reactions were performed using KOD FX Neo with each macrophage genome DNA as a template and the following primer pairs: Forward: 5′-GAGGCATAGTTTCACTCTGTTG-3′ and Reverse: 5′-CTGGACACAGTGGCTCATAC-3′. PCR products were sent to Azenta Japan and complementary sequenced with the following primers: 5′-CAGTGGCTCATACCTGTAACC-3′. Genomic DNA was extracted using DNA lysis buffer (1 M Tris-HCl, pH 8.0, 0.5 M EDTA, 10% SDS, 5 M NaCl, and 100 µg/mL Proteinase K).

### RNA library preparation and RNA-seq

Approximately 200 ng total RNA was used to generate single-end cDNA libraries using TruSeq stranded mRNA Library Prep (Illumina, San Diego, CA). The libraries were sequenced using a NovaSeq 6000 Sequencing System in 100 base single reads and all samples had around 120M reads. Library construction and sequencing were performed at the Genome Information Research Center, Osaka University.

### RNA-seq data processing and DEG analysis

HISAT2 (*55*) was used to perform alignments to the MesAur1.0 golden hamster genome. Transcripts were assembled using StringTie ver 1.3.6 (*56*). Gene and transcript counts were generated with StringTies prepDE.py script. The raw count data from duplicates were analyzed using DEseq2 (*57*) to obtain DEGs between day 0 and day 5. The significance of DEGs with a p-value < 0.1 were shown as a volcano plot using the “EnhancedVolcano” R package (https://github.com/kevinblighe/EnhancedVolcano). A heatmap was generated using the “heatmap.2” R package (https://cran.r-project.org/web/packages/gplots/index.html).

### Enrichment pathway analysis

Extracted DEGs were converted to human homologue genes, hsapiens_associated_gene_name using the “Biomart” R package (*58*). GSEA was inferred using the “fgsea” R package (https://github.com/ctlab/fgsea/) with h.all.v7.5.symbols.gmt for 1000 permutations. Representative pathways were highlighted by the plotEnrichment function.

### Gene ontology

Overall, 5537 upregulated genes were selected and analyzed with g:Profiler (*59*). The GO terms “Cell death”, “Autophagy”, “Immune Response”, “Cytokine Production”, and “Myeloid Leukocyte Activation” were selected to create Venn diagrams of significantly associated genes using VennPlot (http://bioinformatics.psb.ugent.be/webtools/Venn/).

### Splicing QTL analysis

We used the datasets (SNP array and RNA-seq data) of previous eQTL studies obtained from two Europeans cohorts: the DICE (database of immune cell expression, expression quantitative trait loci, and epigenomics) project (*38*) (the database of Genotypes and Phenotypes (dbGaP), phs001703.v1.p1) and the EvoImmunoPop project (*39*) (European Genome-phenome Archive [EGA], EGAS00001001895). We additionally performed genotype imputation using SNP array data. Pre-imputation quality control (QC) of the genotyping data was performed using PLINK 2.0 (https://www.cog-genomics.org/plink/2.0/) with the following parameters (--mind 0.02 --king-cutoff 0.0884 --geno 0.01 --maf 0.01 --hwe 1e-5). Post-QC variants were prephased using SHAPEIT (*60*) and imputation was performed using MiniMac3 (*61*) and 1000 Genomes Phase 3 (release 5) as the reference panel (*62*). Post-imputation QC was performed using PLINK 2.0 with the following parameter (--minimac3-r2-filter 0.3). Genotyped and imputed autosomal SNPs or indels with minor allele frequency (MAF) ≥ 0.01 were used for subsequent QTL analysis with a related expression dataset. We re-aligned the RNA-seq reads on the GRCh38 genome using STAR software v2.7.0 (*63*) in two-pass mode with the GENCODE38 annotations combined with the *de novo* fusion transcripts obtained from long-read capture RNA-sequencing. We conducted junction-based sQTL analysis using LeafCutter (*64*). For the normalization of junction read counts, we performed quantile normalization, rank-transformed normalization, and PEER normalization using 15 hidden factors (*65*). Correlation analysis of the genotype with the junction read count was performed for variants with a MAF ≥ 0.01 within a 1-Mb window around each transcript using the MatrixEQTL R package (*66*) with the top 10 genetic principal components as covariates.

### Colocalization analysis of sQTL and GWAS signals

We used the GWAS summary statistic of severe COVID19 (COVID-19 HGI release5 (https://storage.googleapis.com/covid19-hg-public/20201215/results/20210107/COVID19_HGI_A2_ALL_eur_leave_23andme_20210107.b37.t xt.gz). To evaluate the colocalization of sQTL and GWAS signals, we applied a Bayesian framework using the coloc R package (*67*). We tested for the 500,000 bp window centered on the GWAS lead variant and we considered PP-H4 (posterior probability of shared causal variant) > 0.8 to indicate significant colocalization. Colocalization plots were generated using LocusCompare (*68*).

### Long-read capture RNA-sequencing

We prepared xGen Custom Target Capture Probes (biotinylated 120bp-ssDNAs generated by IDT, Coralville, IA, USA) that covered the entire main-isoform sequences of *IFNAR2* (ENST00000342136.9) and *IL10RB* (ENST00000290200.7) as well as the junction sequences of the novel transcripts. We isolated total RNA from THP-1 cells with or without stimulation by PMA (10 ng/ml for 24 h/72 h), LPS (100 ng/ml for 24 h/72 h) or human IFN-γ (10 ng/ml for 24 h/72 h). We reverse transcribed 100 ng of total RNA by smartseq v2 protocols (*69*) with oligo-dT primers and then amplified them by 22 cycles of PCR using the KAPA HiFi Hot Start Ready Mix (Kapa Biosystems, Wilmington, MA, USA) with 5Me-isodC-TSO and ISPCR primers. We hybridized and captured cDNA with xGen probes using an xGEN Hybridization and Wash Kit (IDT). We then amplified the captured cDNA with an additional 14 cycles of PCR as described above. For library preparation for sequencing, we used a Nanopore Ligation Sequencing Kit (Oxford Nanopore Technologies, Oxford, United Kingdom) and NEBNext Quick Ligation Module/NEBNext Ultra II End-Repair/dA-Tailing Module (New England Biolabs). Then, cDNAs were sequenced by Flongle Flow Cell (FLO-FLG001; Oxford Nanopore Technologies). Basecalling was performed using Guppy (v4.4.1). The obtained fastq files were aligned to the GRCh38 primary assembly using minimap2 v2.17 with reference to the splice junctions in the GENCODE38 annotation. We used the flair pipeline (*70*) to identify the full-length of the novel transcripts and filtered them using the following criteria: 1) isoforms expressing more than 50 reads in total, 2) isoforms whose 5′ end was located within 100 bp from the FANTOM CAGE peak (TSS peak based on a relaxed 0.14 threshold by TSS classifier), 3) isoforms whose 3′ end is located within 100 bp from the TES of PolyASite2.0, and 4) isoforms evaluated as protein coding isoforms by CPAT v3.0.4 (coding probability ≥ 0.364) (*71*).

### Three-dimensional structure computational analysis

Three-dimensional (3D) structures of the hybrid receptor and receptor-ligand complexes were predicted by AlphaFold v2.2.2 (https://github.com/deepmind/alphafold) (*72*). For each receptor-ligand complex, 25 structural models were generated (5 predictions for each of 5 AlphaFold machine learning models). Predicted DockQ (pDockQ) scores (https://gitlab.com/ElofssonLab/FoldDock/-/blob/main/src/pdockq.py) (*73*) were calculated to evaluate the binding affinity of complex models, based on predicted structures and their predicted interface lDDT (plDDT) scores. A violin plot was drawn by Matplotlib (https://ieeexplore.ieee.org/document/4160265) to compare the distribution of pDockQ scores of different complexes. Because AlphaFold modelling is not restricted by the transmembrane domain position, a follow-up step was carried out to construct membrane-embedded complex models. The extracellular domains were first trimmed out of the complex models, and the transmembrane domains were then repositioned onto the same plane beneath the C-terminus of the extracellular domains. They were joined by rebuilding joint regions with the protein structure modelling program MODELLER (*74*). Coarse-grained lipid bilayer models were generated by BUMPy (*75*) for visualization. Structural visualization was performed by PyMOL (The PyMOL Molecular Graphics System, Version 2.5.2, Schrödinger, LLC, New York, NY, USA).

## Statistical analyses

Statistical analyses were performed using GraphPad Prism 8.0. Statistical significance was determined by two-tailed, unpaired parametric *t*-tests or Mann–Whitney *U-*test. For multiple experimental groups, one-way ANOVA with Tukey multiple comparisons was used. Quantitative data are presented as the mean ± standard deviation (SD). Differences were considered significant when *p < 0.05, **p < 0.01, ***p < 0.005, or ****p < 0.001. Additional statistical details are shown in the figure legends.

## Acknowledgments

We thank Y. Sakamaki of the Tokyo Medical and Dental University Research Core for her expert technical assistance with electron microscopy. We also thank Y. Seki and K. Kobayashi for assistance with experiments, M. Kinoshita and Y. Kawada for secretarial assistance, and T. Kojima for technical assistance. We would like to thank all the participants in our institute for the management of patients with COVID-19. We thank J. Ludovic Croxford, PhD, from Edanz (https://jp.edanz.com/ac) for editing a draft of this manuscript. Graphical abstract was created with BioRender.com. This work was supported by Innate Cell Therapy Co., Ltd.

## Funding

This work was supported by the Japan Science and Technology Agency (JST) by funding with a Grant-in-Aid for Transformative Research Areas B (22H05060 and 22H05061) and Grant-in-Aid for Scientific Research on Innovative Areas (18H05032). This work was supported by the Japan Agency for Medical Research and Development (AMED); Research Program Immunology and Allergy under grant number 21ek0410083h0002 and the Research Program on Hepatitis (18fk0310106h0002 and 18fk0210041h0001). This work was also supported by the Takeda Hosho Grants for Research in Medicine and the Visionary Research Fund from the Takeda Science Foundation, JST Moonshot R&D under Grant Number JPMJMS2024, Grant-in-Aids for Scientific Research (B) (22H02597), Grant-in-Aid for Challenging Research (21K19501) from the MEXT of Japan, and a grant from Medical Research Center Initiative for High Depth Omics.

## Author contributions

Y. Mitsui. designed and performed the experiments and wrote the manuscript. T. Suzuki. and T.O. performed the experiments with SARS-CoV-2 and drew the diagrams. J.I. and Y.K. performed the GWAS and QTL analysis. K.Y. and Y.K. performed the long-read capture RNA-seq analysis and computational structural analysis. K.K., T.K., and J.W.S. performed the bioinformatic analysis. M.K., J.W., and M.E. performed the experiments. S.L. and D.M.S. predicted the molecular mechanism of CiDRE using structural modeling. Y. Miyazaki. collected the patient information and samples. S.O., T.H., and S.Y. designed the classification criteria and performed the ELISAs. S.K., N.S., A.K., and S.A. assisted with the experiments. T. Satoh. designed the experiments, wrote the manuscript and supervised the project.

## Competing interests

Authors declare that they have no competing interests.

## Data and materials availability

The RNA-seq data are available in the DDBJ database under BioProject accession number PRJDB14430. This paper does not report original code. Any additional information required to reanalyze the data reported in this paper is available from the lead contact upon reasonable request.

## Supplemental Figures and Figure legends

**Fig. S1.**
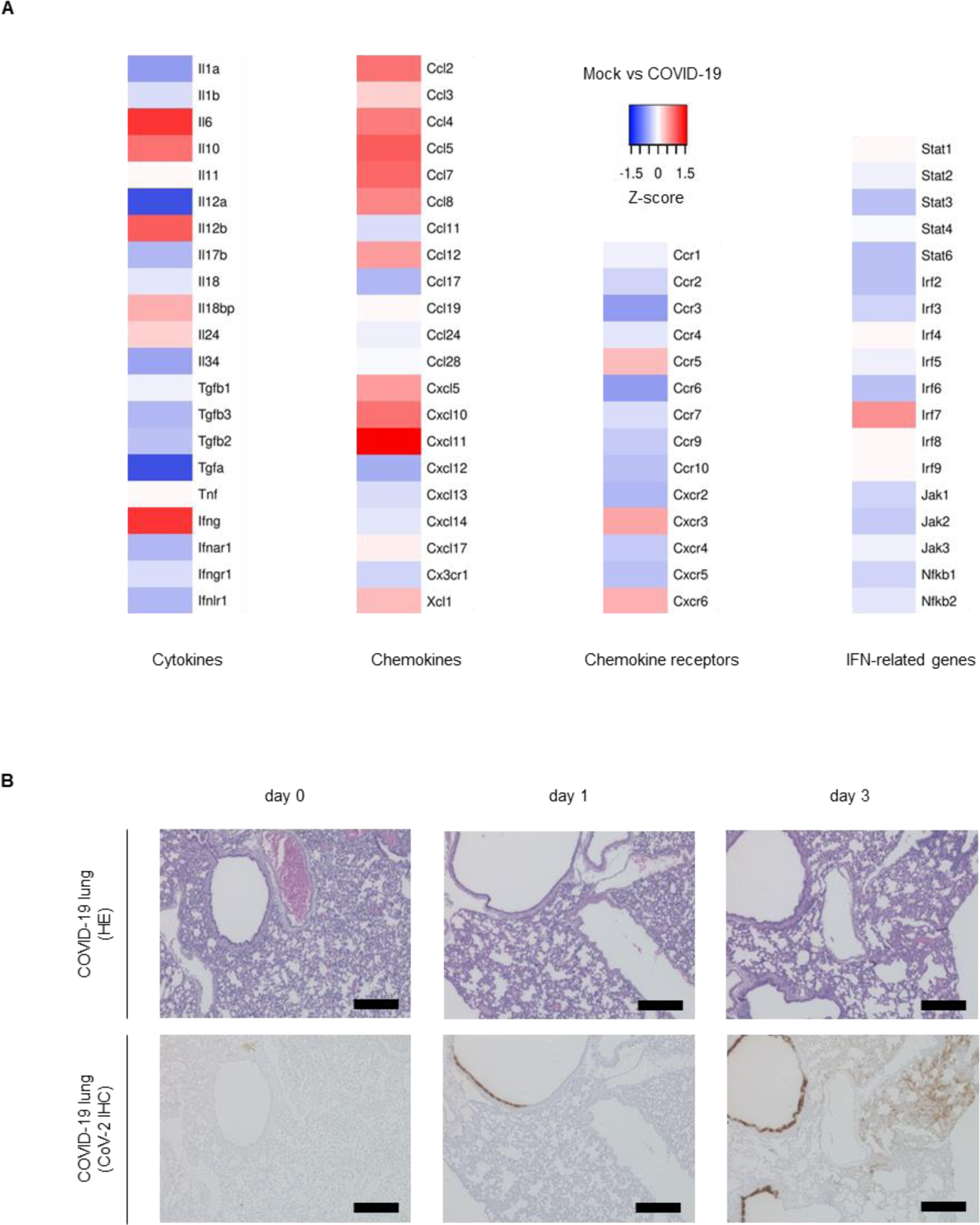
RNA-seq and pathological analysis of the Syrian hamster COVID-19 model. **(A)** Heatmap showing DEGs for genes of interest related to immune and virus-related cellular processes between the Mock and COVID-19 groups (n=2). **(B)** HE and immunohistochemistry of SARS-CoV-2 in the lungs of Syrian hamsters post-infection. The images were focused on the bronchial epithelium and lung parenchyma. Scale bars, 100 µm.

**Fig. S2.**
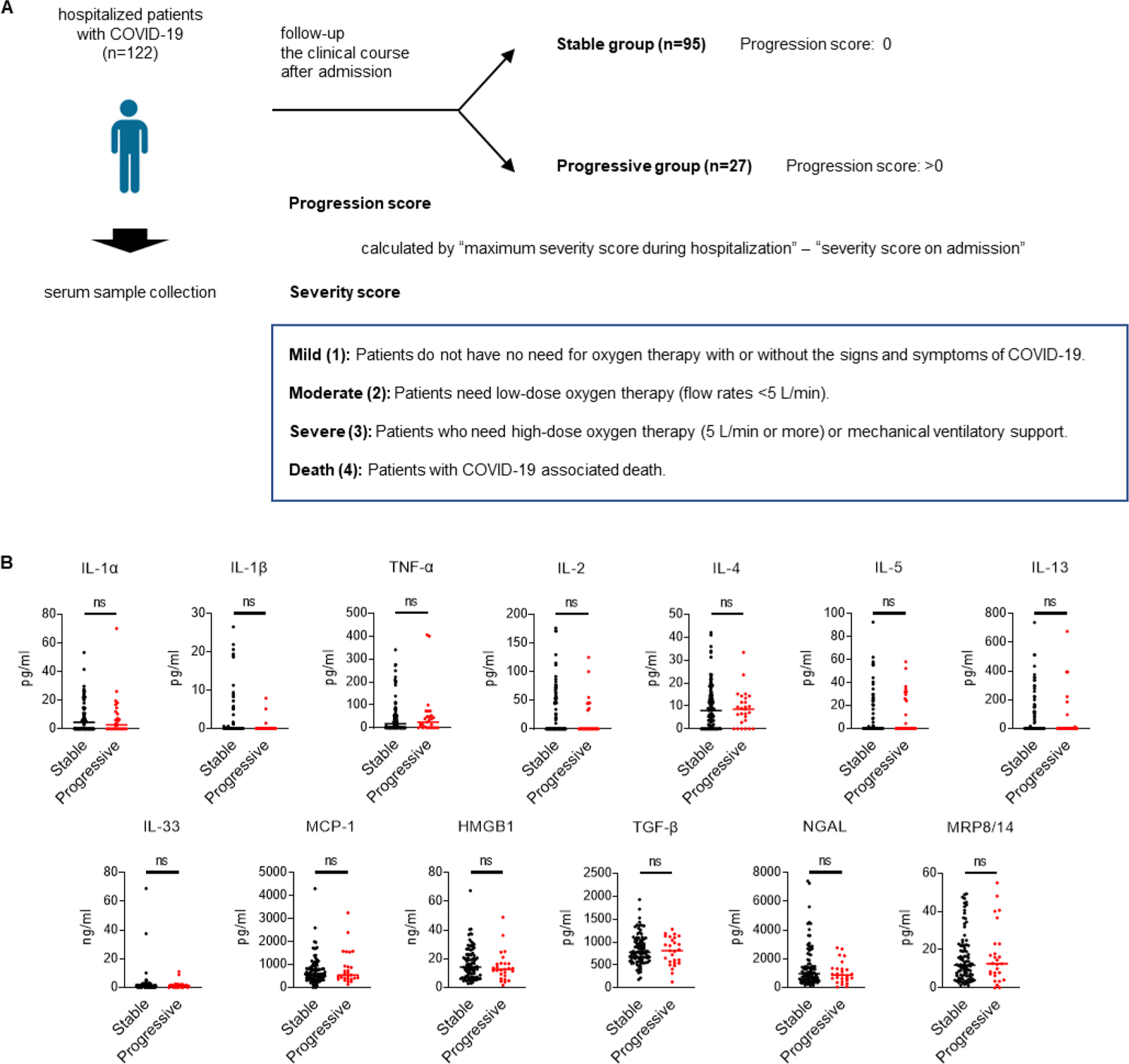
ELISA data sets of human serum. **(A)** Grouping diagram of enrolled patients for serum ELISA. Based on their clinical course, COVID-19 patients were divided into Stable (n=95) or Progressive groups (n = 27) respectively. All patients were classified according to the Japanese clinical guidelines, “Clinical Management of Patients with COVID-19 (Ministry of Health, Labour and Welfare, Japan.)”, as described previously (*76*). Created with BioRender.com. **(B)** Serum levels of several proteins were compared between the Stable (n=95) and Progressive (n=27) groups. The dots represent individual patients and the error bars indicate the median. Statistical significance was determined by a Mann–Whitney *U-*test.

**Fig. S3.**
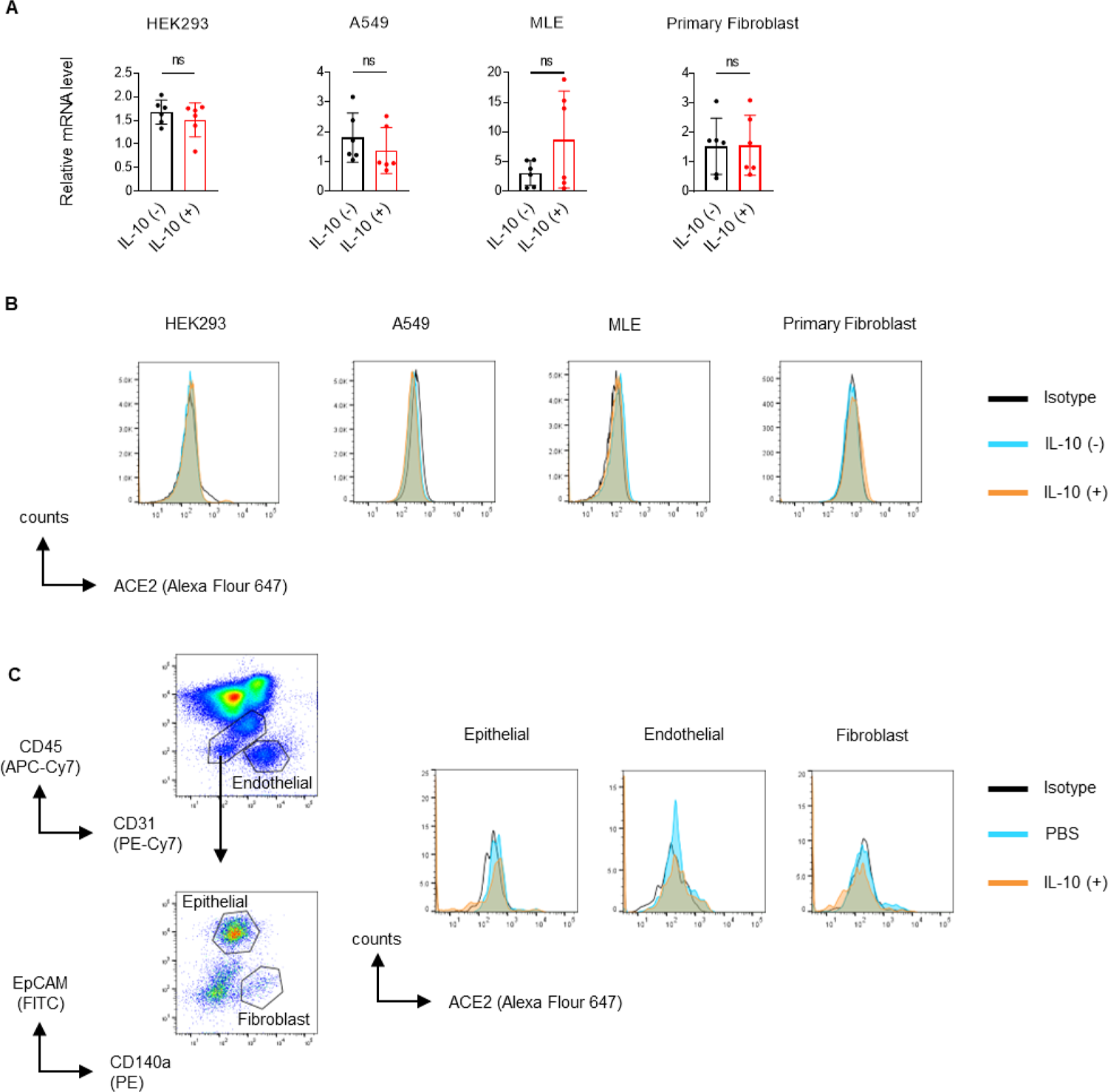
ACE2 expression in epithelial and stromal cells under IL-10 stimulation. **(A)** qPCR and **(B)** flow cytometric analysis of ACE2 expression in human and murine cell lines and murine primary cells. **(C)** Flow cytometric analysis of murine lungs intratracheally injected with IL-10 (10 µg/animal). Gating strategy of each population is shown in the left panel. Samples were collected 3 days after injection and gated as indicated above. The error bars indicate mean ± SD. Statistical significance was determined by two-tailed, unpaired parametric *t*-tests. Data are representative of at least three independent experiments.

**Fig. S4.**
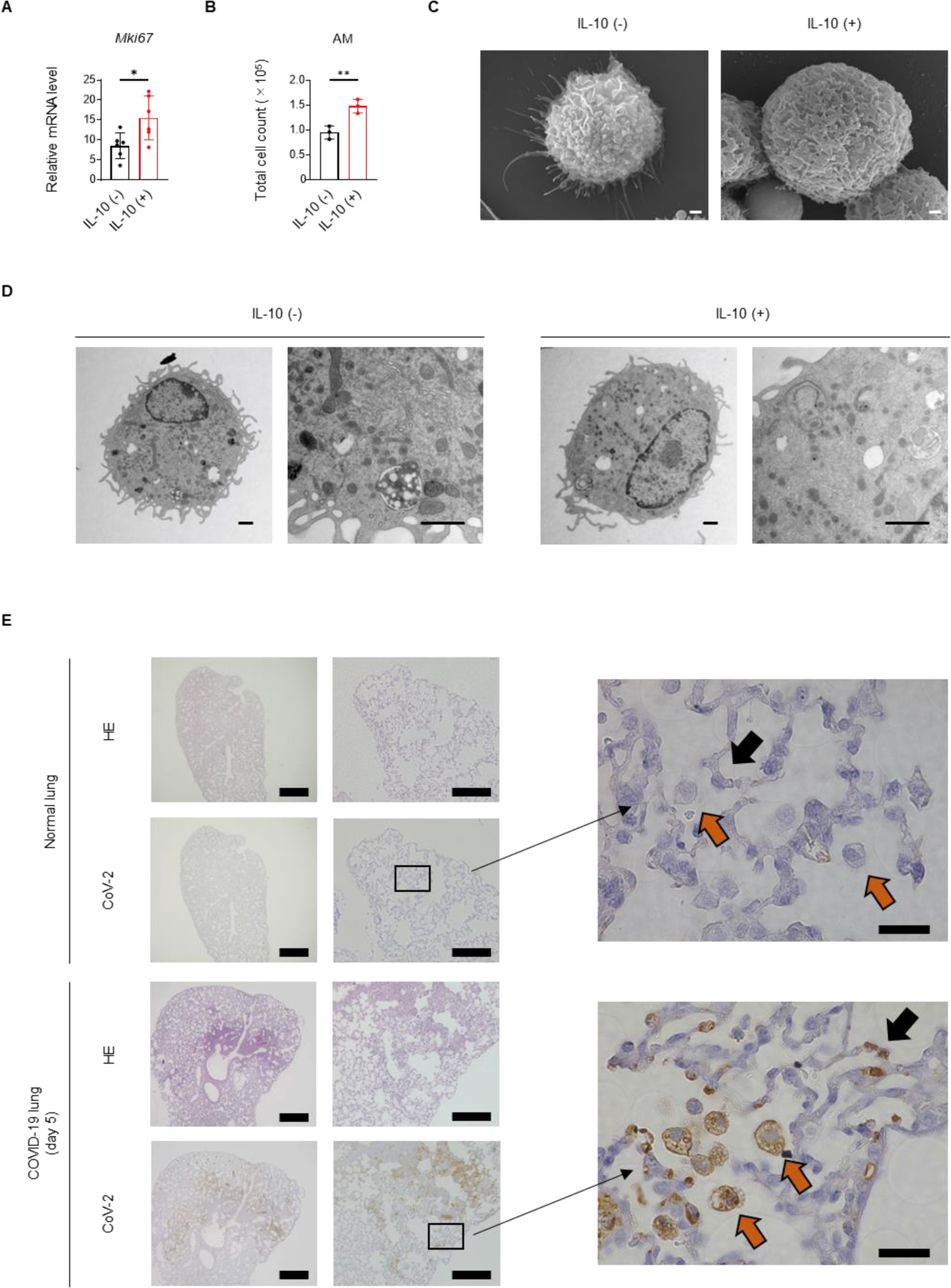
Morphologic and phenotypic changes in alveolar macrophages after IL-10 induction and SARS-CoV-2 challenge. **(A)** The expression of *Mki67* in murine alveolar macrophages stimulated by 40 ng/ml IL-10. **(B)** The number of alveolar macrophages obtained from the bronchoalveolar lavage collected at 3 days after intratracheal administration of IL-10. The error bars indicate mean ± SD. Statistical significance was determined by two-tailed, unpaired parametric *t*-tests. Data are representative of at least three independent experiments. **(C and D)** The morphological changes of alveolar macrophages after IL-10 stimulation (40 ng/ml for 72 h) *in vitro*. The images created by scanning electron microscopy **(C)** and transmission electron microscopy **(D)** were shown. Scale bars, 1 µm. **€** HE and immunohistochemistry of SARS-CoV-2 in the lungs of Syrian hamsters post-infection. The images were focused on the alveoli. Scale bars, 500 µm (left), 100 µm (middle), and 20 µm (right). Orange and black arrows indicate alveolar macrophages and epithelial cells, respectively.

**Fig. S5.**
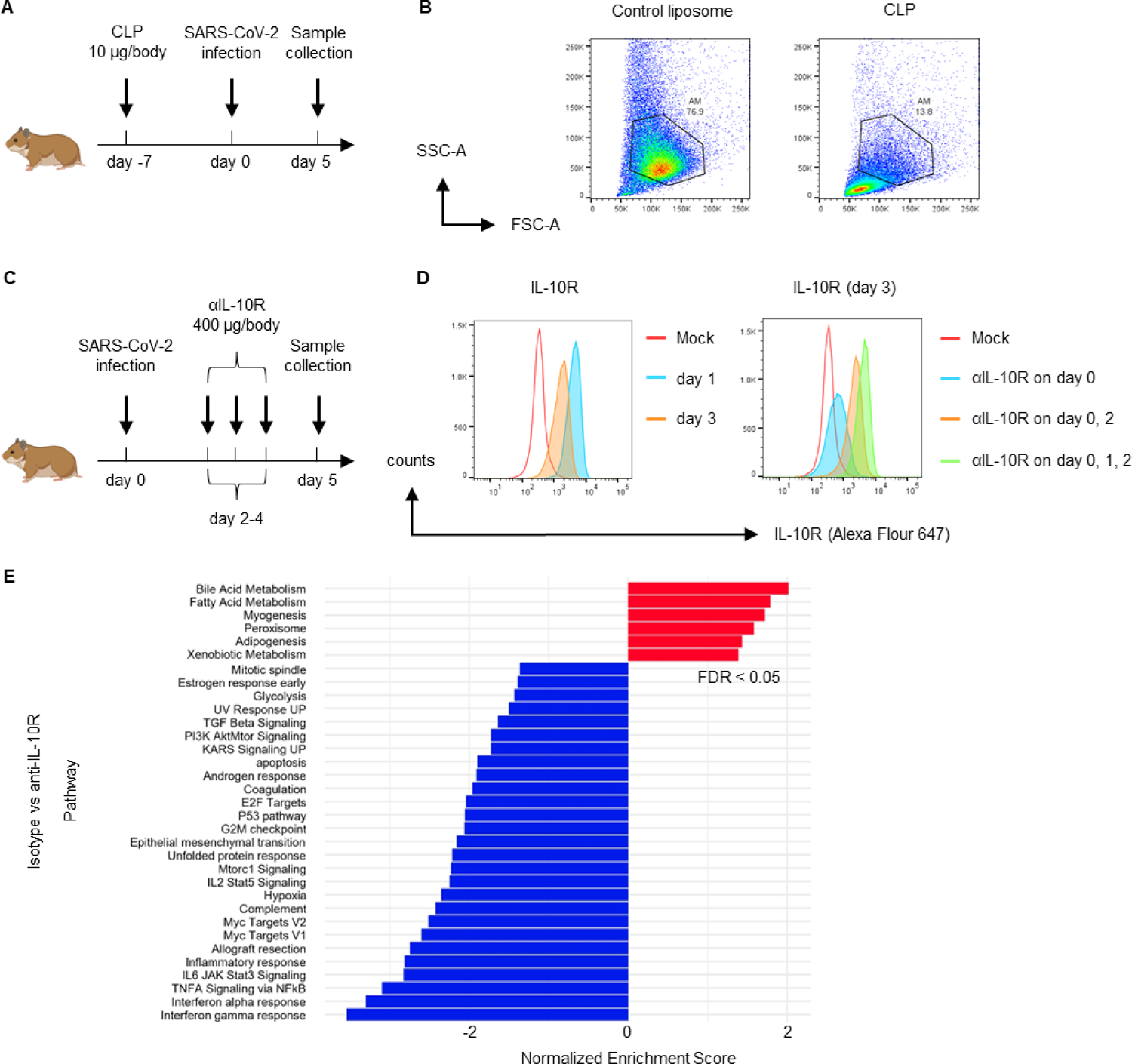
Alveolar macrophage depletion and IL-10R antibody administration model in hamsters. **(A)** Diagram showing the experimental protocol for alveolar macrophage depletion. Created with BioRender.com. **(B)** Flow cytometric analysis of CLP-induced lungs at day 3 after injection. Alveolar macrophages were successfully depleted in this model. **(C)** Diagram showing the experimental protocol for IL-10R antibody administration model. Created with BioRender.com. **(D)** Flow cytometry analysis of IL-10R antibody-induced lungs. After a single administration (left), IL-10R antibody binding on the cell surface of alveolar macrophages was confirmed for 1–2 days. After multiple administration (right), IL-10R antibody binding was enhanced compared to a single administration. **(E)** RNA-seq analysis of the IL-10R antibody administration model. Hallmark plot of GSEA enrichment analysis between the Isotype and anti-IL-10R groups (n=2). Enrichment in the anti-IL-10R group is shown as a positive normalized enrichment score (NES). Statistically significant gene sets with FDR < 0.05 are shown.

**Fig. S6.**
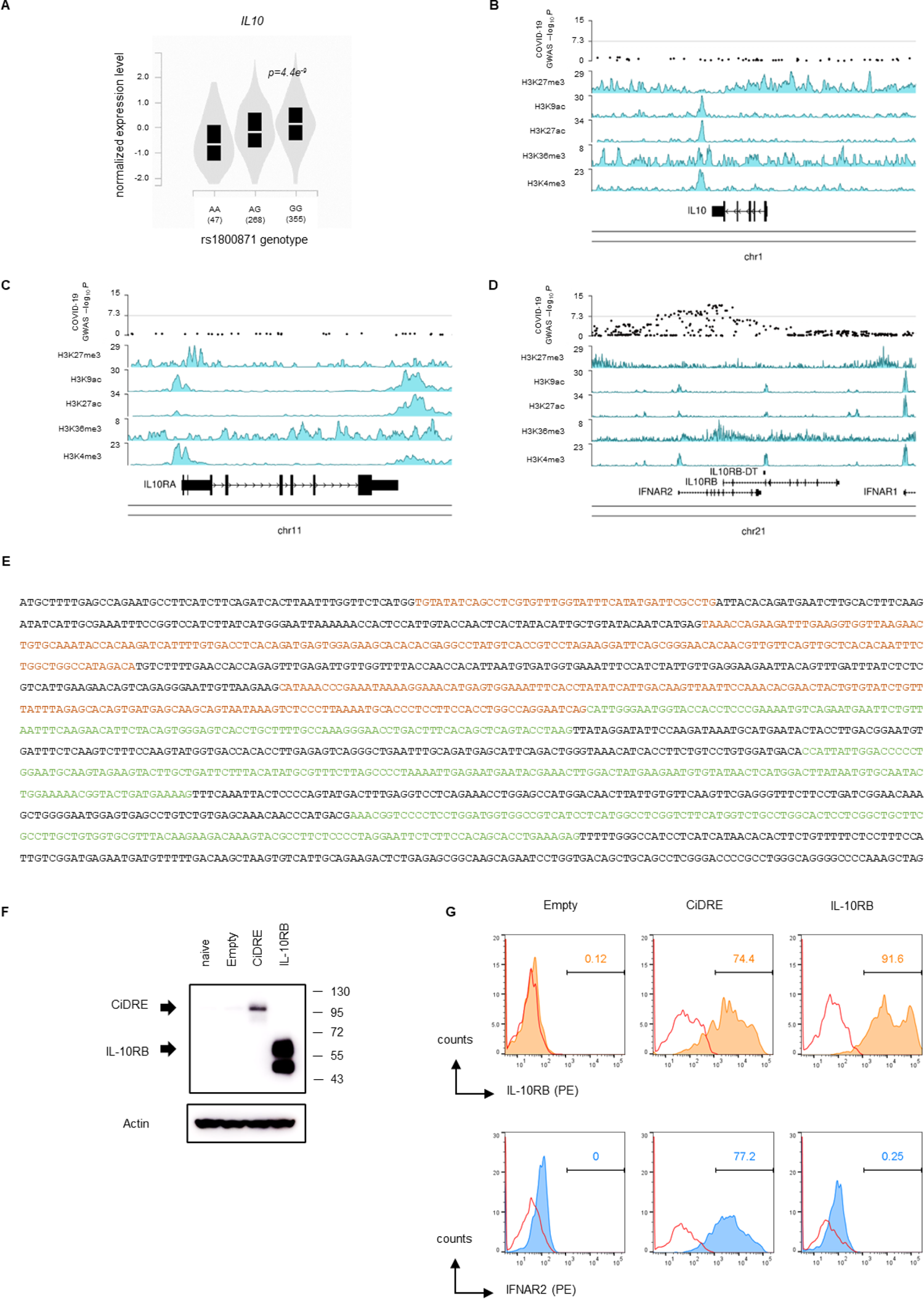
Novel transcripts in the *IFNAR2/IL10RB* lesion were involved in COVID-19 severity as functional mRNA encoding the IFNAR2-IL-10RB hybrid receptor. **(A to D)** Regional associations of the *IL10* and *IL10R* locus in genome-wide association studies of sever COVID-19. **(A)** Violin plot of *IL10* expression showing the eQTL effect based on rs1800871 at the *IL10* promoter locus obtained from the GTEx portal (https://gtexportal.org/). **(B-D)** Manhattan plot showing GWAS results from COVID-19 HGI A2 datasets for COVID-19 severity and histone modification plots at the *IL10* **(B)**, *IL10RA* **(C)** and *IL10RB* **(D)** loci. **(E)** Coding sequence (CDS) of the novel transcripts encoding the IFNAR2-IL-10RB hybrid product. The CDS of the transcripts contains 1638 nucleotides. The CDS consists of the front part of *IFNAR2* (exons 2-7) and the rear part of *IL10RB* (exons 2-7). Individual exons are colored alternatively (*IFNAR2*: orange and black, *IL10RB*: green and black). **(F and G)** Protein expression analysis of the IFNAR2-IL-10RB hybrid product. The IFNAR2-IL-10RB hybrid protein encoded by the novel transcripts was named CiDRE. NanoLuc (empty control), CiDRE, and human IL-10RB overexpressing HEK293 cell lines were used. **(F)** The successful translation of CiDRE was confirmed by western blot analysis using an anti-IL-10RB monoclonal antibody (clone 90220). **(G)** The cell surface expression of CiDRE was confirmed by flow cytometric analysis using anti-IL-10RB and IFNAR2 monoclonal antibodies (clones 90220 and 122).

## Supplementary Material

**Table S1.** Gene ontology analysis of the Syrian hamster COVID-19 model.

**Table S2.** RNA-seq analysis in the Syrian hamster COVID-19 model. Differentially-expressed genes between Mock (day 0) and COVID-19 (day 5) groups were listed.

**Table S3.** Patients characteristics. p values were calculated by parametric t test or Mann-Whitney U test for numerical variables and Fisher’s extract test for two categorical variables. Statistical significance was defined by two-tailed p < 0.05 as indicated with an asterisk. SD: standard deviation. IQR: Interquartile range.

**Table S4.** RNA-seq analysis in IL-10R antibody administration model. Differentially-expressed genes between Isotype (IL10AbMinus) and COVID-19 (IL10AbPlus) groups were listed.

**Table S5.** Long-read capture RNA-seq analysis of *IFNAR2/IL10RB*lesion in the monocyte/macrophage transcriptome. Two transcripts which meet the criteria described in Materials and Methods were listed. Both transcripts contain the same CDS that is translated into CiDRE.

**Table S6.** qPCR primer sequences, reagents, and materials.

## Notes

### Competing Interest Statement

The authors have declared no competing interest.

